# IMPASTO: Multiplexed cyclic imaging without signal removal *via* self-supervised neural unmixing

**DOI:** 10.1101/2022.11.22.517463

**Authors:** Hyunwoo Kim, Seoungbin Bae, Junmo Cho, Hoyeon Nam, Junyoung Seo, Seungjae Han, Euiin Yi, Eunsu Kim, Young-Gyu Yoon, Jae-Byum Chang

## Abstract

Spatially resolved proteomics requires a highly multiplexed imaging modality. Cyclic imaging techniques, which repeat staining, imaging, and signal erasure, have been adopted for this purpose. However, due to tissue distortion, it is challenging to obtain high fluorescent signal intensities and complete signal erasure in thick tissue with cyclic imaging techniques. Here, we propose an “erasureless” cyclic imaging method named IMPASTO. In IMPASTO, specimens are iteratively stained and imaged without signal erasure. Then, images from two consecutive rounds are unmixed to retrieve the images of single proteins through self-supervised machine learning without any prior training. Using IMPASTO, we demonstrate 30-plex imaging from brain slices in 10 rounds, and when used in combination with spectral unmixing, in five rounds. We show that IMPASTO causes negligible tissue distortion and demonstrate 3D multiplexed imaging of brain slices. Further, we show that IMPASTO can shorten the signal removal processes of existing cyclic imaging techniques.

## Introduction

Multiplexed imaging, which enables the visualization of a large number of targets in the same specimen, has received considerable attention due to its potential use in precision medicine and in a deeper understanding of biological mechanisms^1–3^. Fluorescence cyclic imaging techniques are frequently used for such purposes^4^. These techniques achieve high multiplexing capability through iterative staining, imaging, and signal removal. Existing cyclic imaging techniques can be divided into three categories based on the signal removal mechanisms: photobleaching-based, chemical-inactivation-based, and antibody-stripping-based methods. In photobleaching-based methods, the fluorophores of antibodies are bleached by directly illuminating the target specimens with a high intensity light^5,6^. In chemical-inactivation-based methods, the fluorophores of antibodies are deactivated by chemical treatment^7–11^. In contrast, antibody-stripping-based methods separate antigen–antibody binding through chemical treatment^12–14^. Several novel cyclic imaging methods based on these signal removal approaches have been reported ^6–14^, which can be used to observe multiple proteins in a single specimen.

In cyclic imaging, the signal removal processes need to be carefully optimized to satisfy two seemingly contradictory requirements: high fluorescence signal intensities and total signal removal. In photobleaching-based methods, the photobleaching time must be extended to several hours or days to completely bleach the residual signal in deeper areas of thick tissue slices^15^. Previous studies on chemical-inactivation-based techniques adopted hydrogen peroxide, but tissue damage or cell loss occurred over repeated cycles, especially in tissue sections^7,11^. The use of lithium borohydride (LiBH4) reduces such tissue damage^10^. In contrast, proteinase-based antibody-stripping techniques result in epitope loss^11,16^. Recently developed antibody-stripping techniques have mitigated this problem^13,14,17^. These cyclic imaging techniques have mostly been used to acquire two-dimensional images of cultured cells or thin tissue sections due to the need for further optimization of the signal removal processes for thick specimens or to the potential distortion of thick specimens^18^ (see **Supplementary Figure 1** for the distortion of thick tissues observed in cyclic imaging). Instead, cyclic 3D imaging has been demonstrated with specialized specimen treatments that greatly enhance the mechanical stability of specimens, including strong glutaraldehyde fixation or hydrogel embedding^19–22^. Recently, a new class of cyclic imaging techniques, which use oligonucleotide-labeled antibodies, has attracted interest due to their gentle chemical processes^23–25^. However, these techniques are not compatible with commercial antibodies and require the optimization of antibody-oligonucleotide conjugation. Moreover, their use for heavily fixed specimens with reduced epitope retention is limited^4^.

In this work, we propose a new cyclic imaging technique called IMPASTO (**IM**aging based on neural unmixing and **P**yr**A**midal **ST**aining of bi**O**molecules), which replaces chemical or optical signal removal processes with a computational unmixing algorithm. In IMPASTO, specimens are iteratively stained with fluorophore-conjugated antibodies and imaged. A stack of resulting images resembles a pyramid, as the *N*+1^th^-round image is a mixed image of the *N*^th^-round image showing *N* proteins and the signals of the *N*+1^th^ protein. (**Figure 1a**). A pair of images from two consecutive rounds is then unmixed using a self-supervised neural unmixing algorithm to acquire an image of the *N*+1_th_ protein (**Figure 1b**; see **Supplementary Figure 2** for the process of acquiring 5-plex images using IMPASTO). We first show that IMPASTO’s neural unmixing algorithm, which does not require any training data other than the input image to be unmixed, can accurately unmix *N* images containing the signals of *N* proteins. We also show that no epitope loss or tissue distortion occur with IMPASTO, as specimens are treated with only a blocking buffer. Then, we demonstrate 30-plex imaging of mouse brain slices in 10 staining and imaging rounds. In IMPASTO, fluorophores are not bleached; hence, any fluorophores can be employed. The *N*-color unmixing capabilities of the neural unmixing algorithm and the compatibility of IMPASTO with a wide range of fluorophores permit the use of diverse spectrally overlapping fluorophores and unmixing of their signals. Using the neural unmixing algorithm, we conduct unmixing within and between rounds and demonstrate 30-plex imaging in only 5 rounds. More importantly, IMPASTO’s ability to preserve tissue integrity enables the volumetric cyclic imaging of mouse brain slices. Finally, we show that IMPASTO can be combined with existing cyclic imaging techniques to achieve even higher multiplexing capabilities or to shorten their signal removal processes.

**Figure 1.**
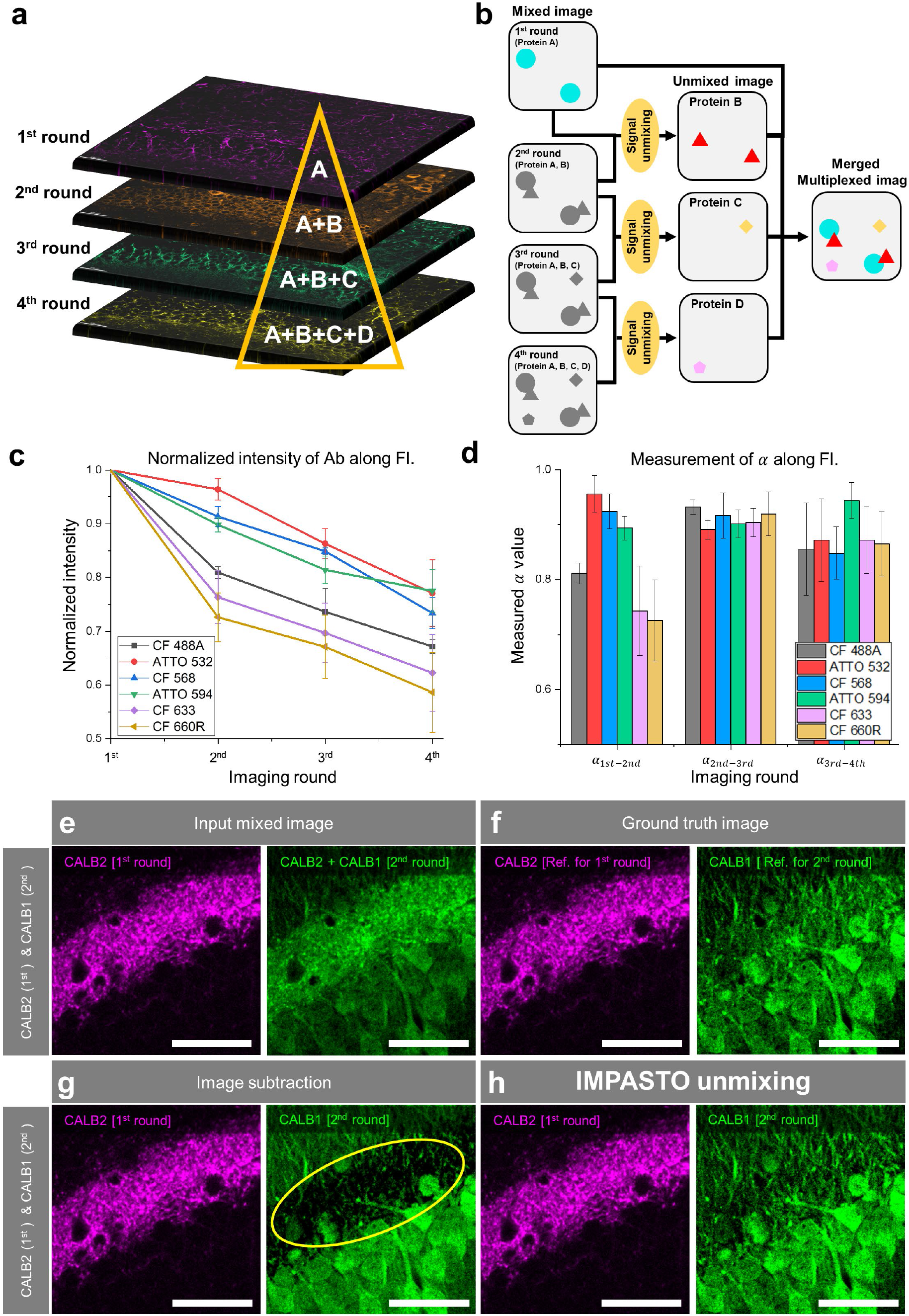
General working principle of IMPASTO. (**a**) Schematic diagram showing how mixed images are acquired in the IMPASTO process. A, B, C, and D are target proteins stained in the first, second, third, and fourth rounds, respectively. No signal removal process is performed between rounds. (**b**) Schematic diagram showing how images of single proteins are retrieved by unmixing images of two consecutive rounds *via* self-supervised neural unmixing. (**c**) Intensity trends across imaging rounds for six different fluorophores. (**d**) Variation of α. α was calculated from the intensity trends shown in **c**. Equation for α calculation is as follows: 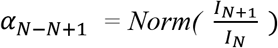, where Norm(I) is normalized intensity, N = 1–3 is the round number. (**e**–**h**) Unmixing of the images acquired from two consecutive rounds using measured α and our neural unmixing code. 50-μm-thick mouse brain slice was immunostained against CALB2 in the first round and against CALB1 in the second round, using the same fluorophore. Note that these two proteins spatially overlap. Simultaneously, these two proteins were immunostained with spectrally distinctive fluorophores, and their images were used as ground-truth images. (**e**) Images acquired in the first and second rounds. Note that the second-round image shows both CALB2 and CALB1. (**f**) Ground-truth images of CALB2 and CALB1. (**g)** Results of the unmixing of the two images shown in **e** using α shown in **d**. Note that the first-round image (=CALB2) was over-subtracted from the second-round image (=CALB1; yellow circle). (**h**) Results of the unmixing of the two images shown in **e** using our neural unmixing algorithm. Note that the unmixed images are identical to the ground-truth images shown in **f**. Scale bar = 50 μm.

## Results

### General working principle of IMPASTO

In IMPASTO, the *N*^th^-round image contains the immunofluorescence (IF) signals of *N* proteins stained during *N* rounds. After staining the specimen with an antibody against the *N*+1^th^ target protein, the *N*+1^th^-round image is acquired. This *N*+1^th^-round image contains the IF signals of *N*+1 proteins. We then obtain the image of the *N*+1^th^ protein by subtracting the *N*^th^-round image from the *N*+1^th^-round image. When *N* is equal to 1, this process can be mathematically expressed as follows:

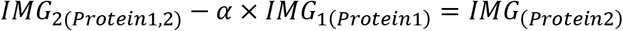

where *IMG*_2(*Protein*1,2)_ is the second-round image showing both the first and second proteins, *IMG*_1(*Protein*1)_ is the first-round image showing only the first protein, *IMG*_(*Protein*2)_ is the image of the second protein, and *α* is a constant (i.e., the value indicating how much of the previous round’s image should be deducted from the current-round image). If *α* can be accurately estimated or measured from separate specimens, the image of the second protein can be retrieved using the formula above. To measure *α* from separate specimens, we stained mouse brain slices with a primary antibody against lamin B1 and a secondary antibody bearing one of six fluorophores (CF 488A, ATTO 532, CF 568, ATTO 594, CF 633, and CF 660R) in the first round. After acquiring the first-round images, we incubated the specimens in a blocking buffer with no antibodies and acquired their second-round images from the same locations. We repeated this process for four rounds with the same imaging conditions and measured the fluorescence signal intensities. As shown in **Figure 1c**, the fluorescence intensities decreased as more imaging rounds were performed due to photobleaching. However, the fluorophores showed different intensity trends due to the difference in their photostability. Even for the same fluorophore, the degree of fluorescence intensity decrease fluctuated across the staining rounds. Such variations in intensity decrease resulted in inaccurate estimations of *α* (see **Figure 1d** for the plot of *α* for each fluorophore across staining rounds). The large standard deviations shown in **Figure 1d** indicate that *α* varied between experiments, even when the same fluorophores were imaged using the same imaging conditions.

The accuracy of the *α* estimation is especially critical for imaging spatially overlapping proteins. If *α* larger than the ground-truth value is used, *IMG*_1(*Protein*1)_ is over-subtracted from *IMG*_2(*Protein* 1,2)_, and the expression level of the second protein is underestimated. If *α* smaller than the ground-truth value is used, *IMG*_1(*Protein*1)_ is under-subtracted from *IMG*_2(*Protein* 1,2)_, and the expression level of the second protein is overestimated. **Figures 1e–h** show such an example. A mouse brain slice was stained with an antibody against CALB2 in the first round and against CALB1 in the second round (**Figure 1e**). These two proteins, which are both calcium-binding proteins and are widely used as cell-type markers in neuroscience, are strongly expressed in the dentate gyrus (DG) region of the hippocampus. We simultaneously stained this specimen with the same antibodies bearing spectrally distinct fluorophores to acquire ground-truth images of CALB1 and CALB2 (**Figure 1f**). When we acquired the image of CALB1 using the predetermined *α* shown in **Figure 1d**, CALB2 was over-subtracted from CALB1, and the resulting image showed a significant signal loss of CALB1 in the inner molecular layer of DG (**Figure 1g**). However, when we estimated *α* using our neural unmixing algorithm, the CALB1 image was successfully retrieved (**Figure 1h**; see **Supplementary Figure 3** for additional examples).

### Neural unmixing: A self-supervised machine learning algorithm for blind signal unmixing

For this unmixing, we devised a self-supervised machine-learning algorithm to unmix *N* images containing the fluorescent signals of *N* proteins without any prior knowledge of the mixing profile (**Figure 2a**). In brief, our neural unmixing algorithm is based on a neural network for independence evaluation.

**Figure 2.**
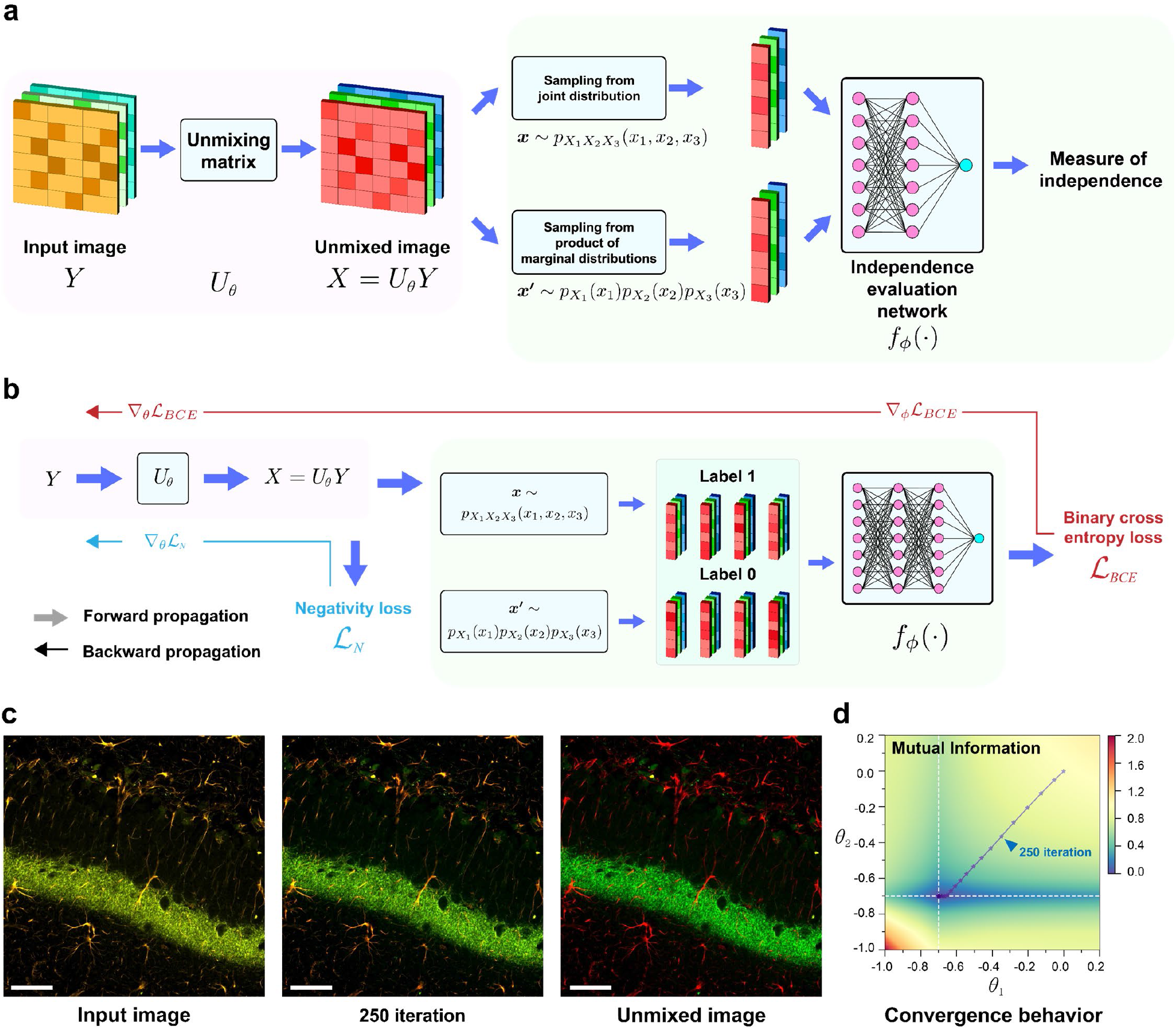
Neural unmixing for the separation of fluorescent signals with no prior knowledge of the mixing profiles. (**a**) The neural unmixing algorithm consists of three components: an unmixing layer, two sampling functions, and an independence evaluation network. The input image first goes through the unmixing matrix, and the pixels are then sampled from the joint distribution (i.e., pixel values from different channels jointly sampled from the same pixel location) and the product of the marginal distribution (i.e., pixel values from different channels sampled independently). Thereafter, two groups of samples are fed into the independence evaluation network to estimate the level of independence. (**b**) The independence evaluation network is trained to classify the samples from the joint distribution and the product of the marginal distribution, whereas the unmixing matrix is trained to make two groups of samples indistinguishable, thereby enforcing independence among different channels of unmixed images. We also positively constrain fluorescent images by employing a non-negativity loss function. (**c**) A synthetic input image obtained by mixing two channel images, an intermediate unmixed image after 250 iterations, and an unmixed image after 1,000 iterations. (**d**) An illustration of changes in mutual information between two channels across iterations. Enforcing independence between two channels minimizes the mutual information between the channels, leading to the parameters of the unmixing matrix converging to the ground-truth values (white dotted line). The unmixing parameters are marked with an asterisk (*) for every 50 iterations. Scale bar = 50 μm.

The neural unmixing consists of three components: an unmixing layer, two sampling functions, and an independence evaluation network. The unmixing layer performs bias subtraction and unmixing as follows:

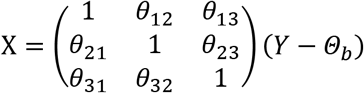

where *θ_ij_*s are the unmixing parameters, *Θ_b_* is the bias parameter, 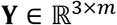 is the three-channel input image with *m* pixels, and 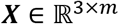 is the three-channel unmixed image with *m* pixels.

Thereafter, two sets of pixels are sampled from the output of the unmixing layer. The first set ***x*** is sampled from the joint distribution (i.e., pixel values from different channels jointly sampled from the same pixel location, 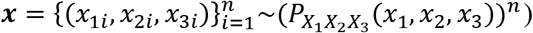, and the second set ***x**′* is sampled from the product of the marginal distributions (i.e., pixel values from different channels sampled independently, 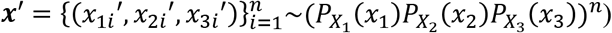 where *n* is the number of samples in each set; *X_k_* is a random variable that corresponds to the sampled pixel value of the *k* -th channel; 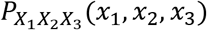 is the joint probability distribution of *X*_1_, *X*_2_, and *X*_3_; 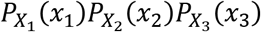 is the product of the marginal distributions; and *x_ki_* is the *k*-th channel pixel value of the i-th sample. One and zero labels are assigned to the first and second sets, respectively. Subsequently, each set of *n* samples is passed to the independence evaluation network.

With such a configuration, the independence among the different channels of the unmixed images is assessed by the independence evaluation network in a way that allows backpropagation for the gradient calculation. Thereafter, the unmixing layer and the independence evaluation network are trained through gradient-based updates in an adversarial manner (**Figure 2b**); the independence evaluation network is trained to classify the samples (i.e., minimize binary cross-entropy loss) from the joint distribution and the product of the marginal distributions, whereas the unmixing layer is trained to make two groups of samples indistinguishable (i.e., maximize binary cross-entropy loss), thereby enforcing independence among different channels of the unmixed images. Consequently, the signals in the two output channels are gradually unmixed, as verified by the decrease in mutual information between each pair of channels across iterations (**Figure 2c,d**).

For two-channel unmixing, we found that the mutual information neural estimator^26^, which estimates the mutual information of two random variables, could be employed as the independence evaluation network instead of the classifier network (see **Supplementary Figure 4**). After the unmixing layer and the independence evaluation network converged, the output of the unmixing layer was taken as the unmixed image.

The major advantage of neural unmixing over existing blind unmixing methods is that any prior knowledge of the unmixed image can be directly incorporated into the optimization problem as a loss function. For example, fluorescent images are positively constrained in neural unmixing by employing a negativity loss function (**Figure 2b**). Moreover, the constant bias in the input image can be estimated and removed as part of the optimization process in neural unmixing.

### Quantitative and qualitative validation of IMPASTO

Next, we quantitatively measured the unmixing accuracy of IMPASTO. We stained mouse brain slices for three rounds, and the target proteins were PV in the first round, S100B in the second round, and GM130 in the third round. We used CF 488A to stain these three proteins for three rounds. To validate the unmixing of IMPASTO, we simultaneously labeled S100B and GM130 with CF 568 and CF 633, respectively, and used these images as the ground-truth images of these two proteins. We included DAPI in all rounds and used it as a registration marker. As shown in the second row of **Figure 3a**, the 488-nm channel image in the first round showed only PV, but the second-round image showed mixed expressions of both PV and S100B, and the third-round image showed mixed expressions of PV, S100B, and GM130. We then acquired an S100B image by unmixing the first- and second-round images and a GM130 image by unmixing the second- and third-round images using the IMPASTO neural unmixing algorithm (**Figure 3b**). The resulting S100B and GM130 images were identical to their ground-truth images (**Figure 3c**; see **Supplementary Figure 5a** for images before and after unmixing and **Supplementary Figure 5b** for additional validation results with the same antibodies). We performed the same validation experiment with different antibodies: lamin B1 in the first round, GFAP in the second round, and CALB2 in the third round. Again, the unmixed images of GFAP and CALB2 matched their ground-truth images (**Supplementary Figure 6**). We measured Pearson correlation coefficients between unmixed and ground-truth images to quantitatively measure the accuracy of neural unmixing. The average was 0.97, which showed that the neural unmixing precisely unmixed *N*^th^ and *N*+1^th^ images to retrieve the image of the *N*+1^th^ protein (**Figure 3d**). We then validated the unmixing accuracy in each round of 10-round staining. In this experiment, we used a 557-nm channel to acquire mixed images and 488-nm and 640-nm channels to acquire ground-truth images. Since ground-truth images of only two proteins could be acquired from one specimen, we performed the 10-round staining and imaging with 5 brain slices, each of which had 2 ground-truth channels. Specifically, we stained 5 brain slices with 10 antibodies bearing CF 568 in 10 rounds. We also simultaneously stained each brain slice with 2 of 10 antibodies bearing CF 488A and CF 633 and then acquired the ground-truth images. For each brain slice, we qualitatively compared the unmixed images with the respective ground-truth images (see **Supplementary Figure 7** for qualitative validation results). The unmixed images showed a high degree of matching with their ground-truth images.

**Figure 3.**
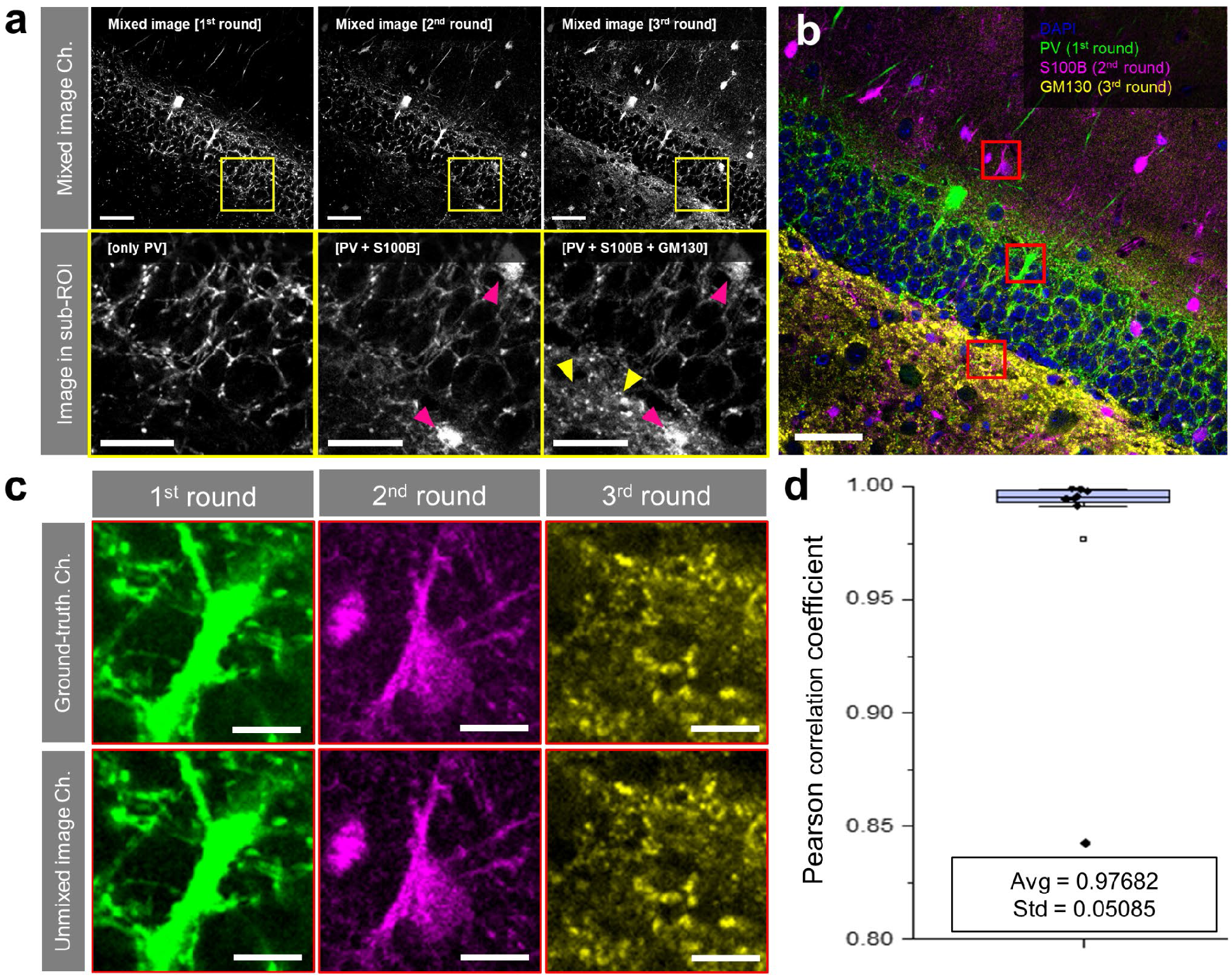
Experimental validation of unmixing of IMPASTO. (**a**) Images of three-round cyclic images obtained without any signal removal process. The three target proteins were simultaneously stained with spectrally distinctive fluorophores, and their images were used as ground-truth images. Top row: mixed images acquired in the first, second, and third rounds. Bottom row: magnified view of the images shown in the top row. Magenta arrowhead: S100B. Yellow arrowhead: GM130. Note that the second-round image contains both PV and S100B, and the third-round image contains PV, S100B, and GM130. (**b**) Overlay of the three proteins after unmixing the mixed images shown in **a** using the neural unmixing algorithm. (**c**) Magnified view of the three red-boxed regions in **b**. Top row: ground-truth images of the three proteins. Bottom row: unmixed images. (**d**) Average Pearson correlation coefficient (r) between the unmixed and ground-truth images measured from 4 specimens (8 images). Scale bar = 50 μm in the first row of (**a**) and in (**b**), 30 μm in the second row of (**a**) and 10 μm in (**c**).

### Demonstration of IMPASTO for 30-plex imaging in 10 rounds

After the thorough validation of our neural unmixing algorithm, we attempted to use 3 colors in each of 10 staining rounds to perform 30-plex imaging. In cyclic staining based on fluorophore bleaching, the complete bleaching of fluorophores needs to be checked after bleaching to ensure that none of the previous fluorescence signals remain. Additionally, fluorophores with high photostabilities, such as Alexa Fluor 546, Alexa Fluor 568, Alexa Fluor 594, and eF615, are not recommended, despite their high brightness and photostability, as they require a longer bleaching process^8,10,28^. However, in IMPASTO, regardless of the degree of photostability, any fluorophores can be used. We used three fluorophores that showed high brightness and photostability in previous studies: CF 488A, CF 568, and CF 633^27^. As shown in **Figure 4**, we successfully demonstrated 30-plex imaging by performing 10 rounds of 3-color staining and imaging using IMPASTO in the DG region of mouse brain slice. We compared the expression patterns of each protein acquired by IMPASTO to the singly stained specimens and found no noticeable differences (see **Supplementary Figure 8** for images of the singly stained specimens, **Supplementary Figure 9** for individual proteins, **Supplementary Figure 10** for the cumulative visualization of the 30 proteins, **Supplementary Video 1** for both individual proteins and cumulative view of all proteins). We successfully demonstrated another 30-plex imaging with different antibody panels (**Supplementary Figure 11** and **Supplementary Video 2**).

**Figure 4.**
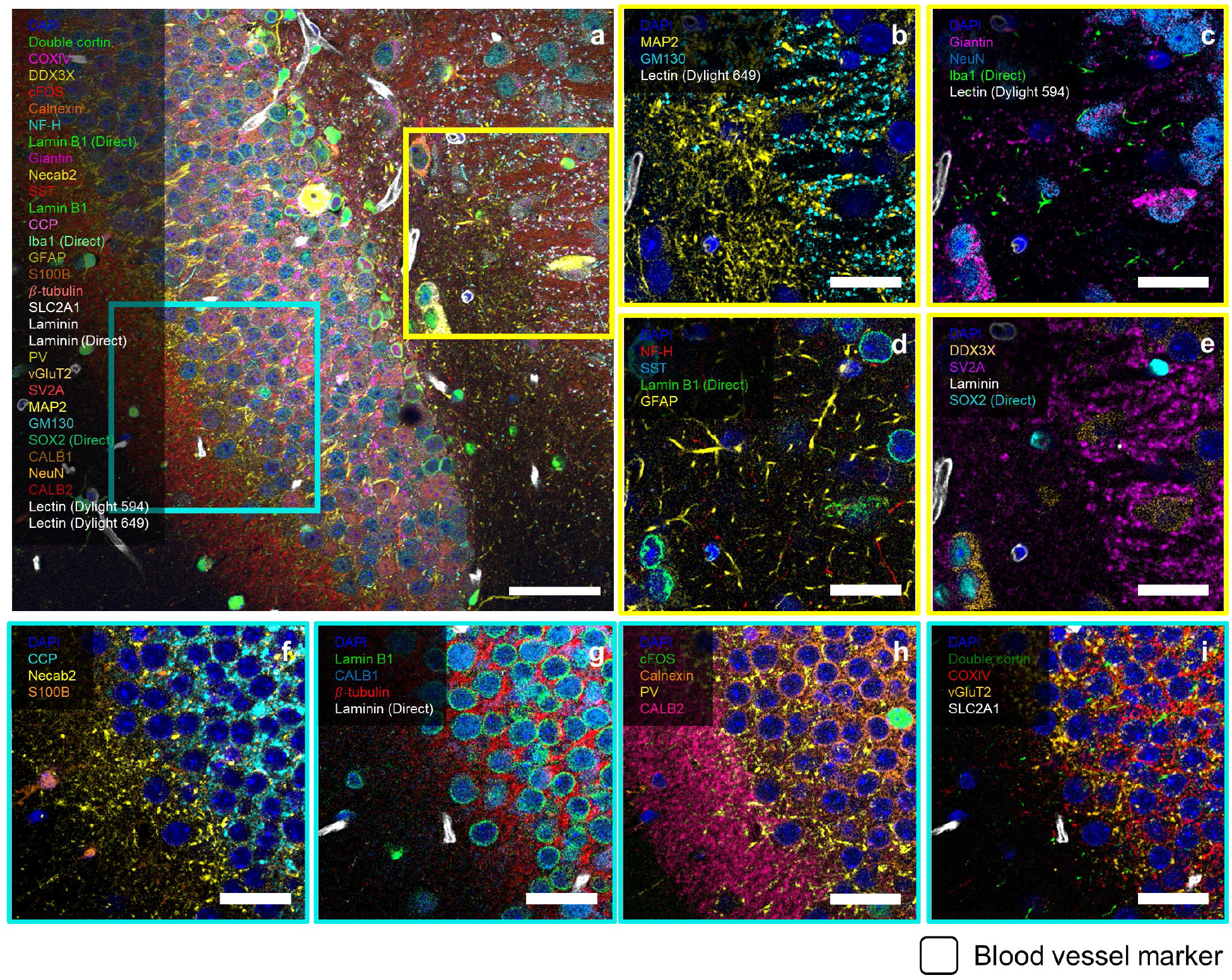
Thirty-plex protein imaging of a mouse brain slice using IMPASTO. Ten rounds of three-plex imaging were performed using IMPASTO in the dentate gyrus of the hippocampal region of a 50-μm- thick mouse brain slice. 488-, 557-, and 640-nm excitation lasers were used. DAPI staining was used as a fiducial marker. (**a**) Merged image of the 30 target proteins. (**b**–**e**) Magnified view of the yellow-boxed region in **a**. For better visibility, four or five proteins are shown in each image. (**f**–**i**) Magnified view of the cyan-boxed region in **a**. Again, for better visibility, four or five proteins are shown in each image. Scale bar = 50 μm in (**a**) and 20 μm in (**b**–**i**).

### Twelve-plex three-dimensional (3D) imaging via IMPASTO

Visualization of numerous protein expressions in a 3D context is essential for obtaining a deeper understanding of the structural information of target tissues^29,30^. The biggest challenge in using cyclic staining for the volumetric imaging of proteins is specimen damage or distortion. To study the origin of tissue distortion, three 50-μm-thick brain slices were first stained with DAPI and then processed with three different buffer exchange procedures for 10 rounds. The first slice was incubated in a wash buffer (1× PBS, 0.2% Triton X-100) for 10 min and then washed three times with the same buffer for 3 min each. The second slice was incubated in a fluorophore-inactivation buffer^10^ for 10 min and then washed three times with a wash buffer for 3 min each. The third slice was incubated in a commercial antibody-stripping buffer for 10 min and then washed three times with a wash buffer for 3 min each. These processes were repeated for these three slices 10 times. To study the distortion of the specimens, the DAPI images were acquired before and after the 10 rounds of buffer exchange. As shown in **Supplementary Figure 1**, the first specimen did not show noticeable distortion after the buffer exchange, indicating that repeated exchange with the same buffer does not mechanically distort specimens. However, the second and third specimens showed distortion both in the axial and lateral directions. Change in the salt concentration or pH induce the swelling or expansion of specimens^31^, which can potentially result in irreversible specimen distortion. IMPASTO uses two buffers: a staining buffer (5% normal serum, 0.2% Triton X-100, 1× PBS) and a wash buffer. As the pH and salt concentrations of these two buffers are almost identical, repeated exchange between them would not distort specimens. Using IMPASTO, we performed 12-plex volumetric imaging with an imaging depth over 20 μm, which is thicker than single cells (**Figure 5**, see **Supplementary Video 3** for a z-stack image and **Supplementary Figure 12** for a magnified view of single-protein 3D images). In volumetric imaging, the fluorescence signal intensities of proteins decreased in the deeper areas. Different proteins showed different intensity decrease trends along the depth of the specimen due to differences in the antibody-binding affinities and abundance of antigens^32^. Even in such cases, IMPASTO successfully unmixed pairs of two consecutive images in all the z-planes shown in **Figure 5**.

**Figure 5.**
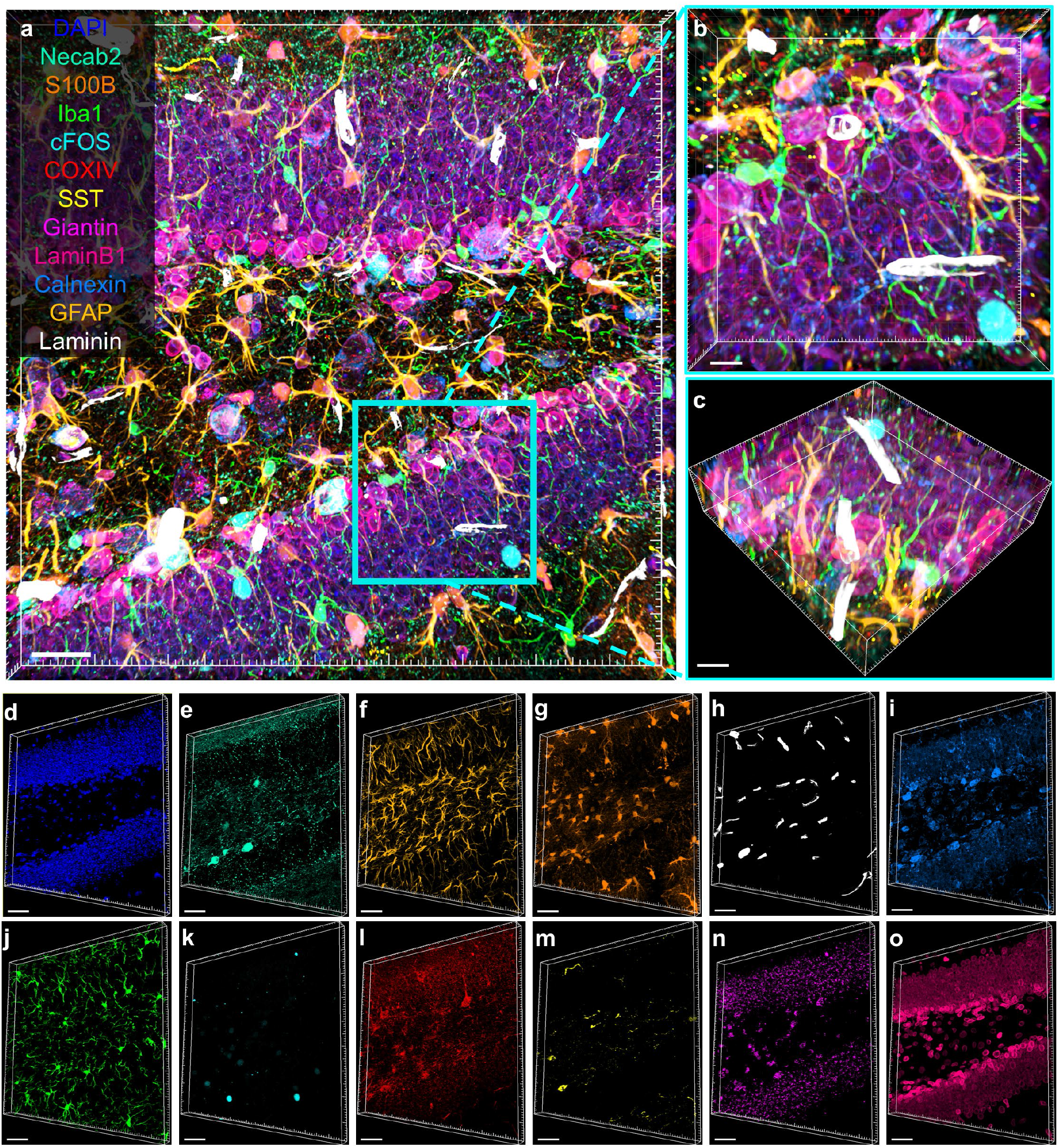
12-plex 3D imaging of a mouse brain slice using IMPASTO. Four rounds of three-plex imaging were performed using IMPASTO in the dentate gyrus of the hippocampal region of a 50-μm-thick mouse brain slice. 488-, 557-, and 640-nm excitation lasers were used. DAPI staining was used as a fiducial marker. In each round, z-stacks were acquired over 20 μm with a step size of 1 μm. Images from two consecutive rounds acquired in the same z-plane were unmixed *via* the neural unmixing algorithm. (**a**) 3D projection view of the merged 12-plex image. (**b**) Magnified 3D projection view of the boxed region in **a**. (**c**) 3D tilted view of the image shown in **b**. (**d**–**o**) Single-protein 3D view. Scale bar = 30 μm in (**a**) and 10 μm in (**b**–**o**).

The mild chemical procedure of IMPASTO would also be beneficial in the imaging of archived clinical formalin-fixed paraffin-embedded (FFPE) specimens. The repeated treatment of FFPE specimens with fluorophore-inactivation buffers^10^ or antibody-stripping buffers could detach the specimens from substrates (see **Supplementary Figure 13a,b** for images of FFPE specimens after 13 rounds of antibody-stripping processes using a commercial stripping buffer and **Supplementary Video 4** for the video of the specimen). However, even after 30 rounds of buffer exchange with only a wash buffer, the FFPE specimen maintained its structure, and no distortion in the staining pattern was observed (**Supplementary Figure 13c,d)**. This result indicates that IMPASTO can be applied to archived clinical FFPE specimens to perform multiplexed imaging without any additional sample treatment process.

### Spectro-temporal unmixing using IMPASTO for enhanced multiplexing capability

While IMPASTO eliminates the need for a signal removal process, it still required more than 10 rounds of repetitive staining and imaging for 30-plex imaging. Since each round required at least several hours of staining, reducing the total number of rounds by using more fluorophores in each round would reduce the total process time and extend the applicability of IMPASTO. To facilitate this, we combined IMPASTO with spectral unmixing. Since our neural unmixing algorithm could unmix two images acquired not only in two separate rounds but also in the same round in different spectral ranges, we used the neural unmixing algorithm for unmixing the signals of spectrally overlapping fluorophores as well as for unmixing the images acquired in two consecutive rounds. Specifically, we used two spectrally overlapping fluorophores for each of the 488-, 557-, and 640-nm lasers and performed six-plex imaging for five rounds. We termed this process spectro-temporal unmixing.

As shown in **Figure 6a–c**, we performed spectro-temporal unmixing, which includes inter-round and intra-round unmixing. In the first round, the first protein was labeled with CF 488A, and the second protein was labeled with ATTO 532. We then acquired two images in two detection channels, each of which included an emission peak of either CF 488A or ATTO 532. We denoted these two images as *IMG*_1,488_ and *IMG*_1,532_. Both *IMG*_1,488_ and *IMG*_1,532_ showed the first and second proteins but with different intensities. In the second round, we labeled the third protein with CF 488A and the fourth protein with ATTO 532. Again, we acquired two images in the same detection channels used in the first round. We denoted these two images as *IMG*_2,488_ and *IMG*_2,532_. Both *IMG*_2,488_ and *IMG*_2,532_ showed the first, second, third, and fourth proteins but with different intensities. To acquire single-protein images of the first, second, third, and fourth proteins from these four mixed images, we first performed inter-round unmixing by subtracting *IMG*_1,488_ from *IMG*_2,488_ and *IMG*_1,532_ from *IMG*_2,532_ using the neural unmixing algorithm (**Figure 6a**). The resulting images, which we denoted *IMG*’_2,488_ and *IMG*’_2,532_, showed the third and fourth proteins with different intensities. We then performed intra-round unmixing (**Figure 6b**). We unmixed *IMG*_1,488_ and *IMG*_1,532_ to retrieve single-protein images of the first and second proteins. Then, we unmixed *IMG*’_2,488_ and *IMG*’_2,532_ to acquire single-protein images of the third and fourth proteins (**Figure 6c**). This spectro-temporal unmixing facilitated enhanced multiplexing, allowing us to perform 30-plex fluorescent imaging in five staining rounds (see **Figure 6d–i** for merged images of each round, **Supplementary Figure 14** for individual proteins, **Supplementary Figure 15** for the cumulative visualization of all 30 proteins, **Supplementary Video 5** for a video showing both individual proteins and the cumulative visualization of all 30 proteins). We successfully demonstrated another 33-plex imaging *via* spectro-temporal unmixing with different antibody panels (**Supplementary Figure 16** and **Supplementary Video 6**). The images acquired in this process had a high enough signal-to-noise ratio to clearly distinguish variations in protein expressions among cells (see **Figure 6j** for six representative cells and **Supplementary Figure 17** for more cells). To validate the cell phenotyping results, we checked whether the previously reported protein expression patterns matched our results. It is known that GFAP- and S100B-positive cells are negative for NeuN^33^; Iba1 expression is not colocalized with NeuN, SOX2, and S100B^34^; S100B expression is not colocalized with NeuN and Iba1^33,35^; and SOX2 is not expressed in granule cells, which are positive for NeuN^36^. All S100B-expressing cells are positive for SOX2, but not all SOX2-expressing cells are positive for S100B^36,37^. All these previously reported findings matched our results, indicating that our proposed spectro-temporal unmixing is suitable for studying the molecular heterogeneity of cells. In our previous work, we showed that blind unmixing could unmix signals of more than 3 fluorophores, and 15-plex imaging could be achieved in a single staining and imaging round^27^. We validated the effectiveness of our neural unmixing algorithm in the spectral unmixing of more than three spectrally overlapping fluorophores through a simulation (see **Supplementary Figures 18** and **19** for the blind spectral unmixing of four and five fluorophores, respectively). The unmixed images were nearly identical to the corresponding ground-truth images with Pearson correlation coefficients higher than 0.98, indicating that the spectro-temporal unmixing proposed herein could be extended to achieve an even higher multiplexing capability.

**Figure 6.**
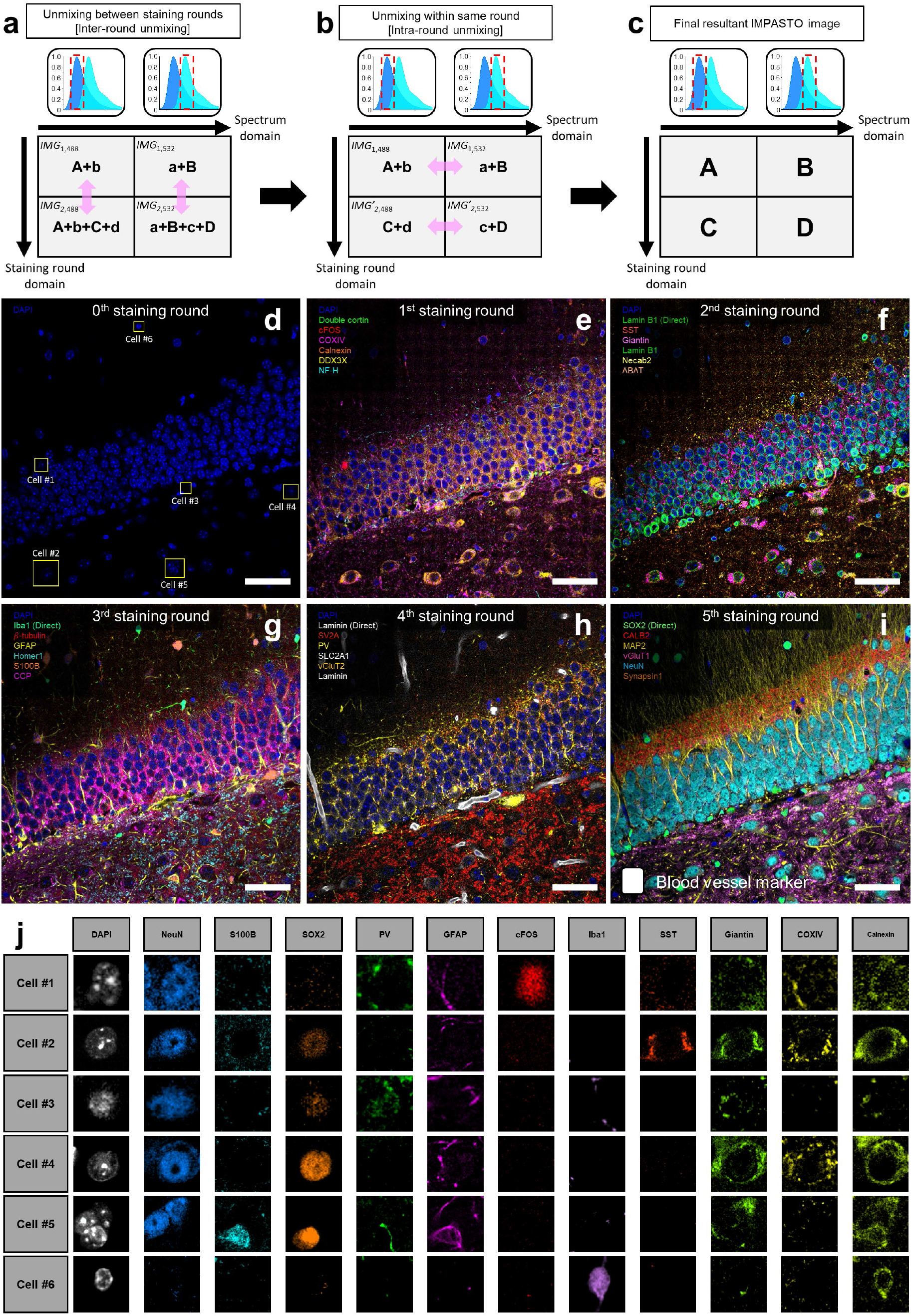
Demonstration of 30-plex protein imaging in 5 rounds using spectro-temporal unmixing. Five rounds of six-plex imaging were performed in the dentate gyrus of the hippocampal region of a 50- μm-thick mouse brain slice. 488-, 557-, and 640-nm excitation lasers were used. DAPI staining was used as a fiducial marker. For each excitation laser, two spectrally overlapping fluorophores were used. In each round, two images were acquired in two different detection ranges, each of which contained the emission peak of the fluorophores. (**a**–**c**) Schematic diagram showing how the spectro-temporal unmixing was conducted. This schematic shows the unmixing procedure for a 488-nm channel. The same procedure was repeated for other channels. Letters (i.e., A, B, C, D, a, b, c, and d) represent stained target proteins. Capital and small letters indicate the relatively stronger and weaker fluorescence intensities of the same protein, respectively. For example, A and a represent high and low fluorescence intensities of the same target protein A. (**a**) Inter-round unmixing. Images from two consecutive rounds were unmixed. (**b**) Intra-round unmixing. Images acquired from the same round but in two different detection ranges were unmixed to separate the signals of the spectrally overlapping fluorophores. (**c**) Unmixed images. (**d**–**i**). Merged images of each round after unmixing. (**j**) Protein expression patterns of six cells shown in the yellow-boxed regions in **d**. Scale bar = 50 μm in (**d–i**).

### Combination of IMPASTO with periodic antibody stripping

Next, we sought to determine the maximum number of rounds that IMPASTO could achieve. IMPASTO uses only mild chemicals (i.e., 5% normal serum, 0.2% Triton X-100, 1× PBS), and similar buffers have been widely used to stain specimens for prolonged periods (longer than a week) without noticeable changes in antigenicity and tissue integrity^22,38–40^. To obtain the 10-round imaging results shown in **Figure 4**, we performed each staining round overnight, and the total staining time was longer than 120 h. Even after such prolonged incubation of the specimens in the blocking buffer, tenth-round staining showed no decrease in antigenicity. In addition, 10 rounds of buffer exchange with brain slices and 30 rounds of buffer exchange with paraffine-embedded specimens did not show any noticeable distortion, as discussed previously. Considering these two observations, epitope loss and tissue distortion were negligible in IMPASTO and therefore would not limit the maximum number of staining rounds that IMPASTO could achieve. We hypothesized that the maximum number of rounds that IMPASTO could repeat would be limited by the steric hindrance of antibodies. Previous studies of cyclic imaging based on fluorophore inactivation reported no steric hindrance, even in 60-plex imaging^6,9^. However, in highly dense biological structures, such as synapses, steric hindrance could limit the multiplexing capability of IMPASTO.

In such cases, we hypothesized that IMPASTO could be combined with antibody-stripping techniques. We performed three rounds of IMPASTO imaging, stripped the bound antibodies from the specimen, and then performed another three rounds of IMPASTO imaging. After the antibody stripping, IMPASTO successfully achieved three-color imaging, which shows that IMPASTO can be combined with antibody-stripping techniques to resolve the steric hindrance issue (see **Supplementary Figure 20**). Interestingly, even when the antibody-stripping process did not completely remove the fluorescent signals, IMPASTO successfully achieved three-color imaging, without being affected by residual signal issues (see **Supplementary Figure 21**). In detail, we performed three-round IMPASTO staining and imaging, followed by the antibody-stripping process. We imaged the antibody-stripped specimen and found that the residual signals remained. We then stained the specimen with the fourth antibody and imaged it. The fourth-round image contained both the residual signals and the signal of the fourth protein. By unmixing these two images, we successfully acquired an image of the fourth protein. We proceeded to the fifth- and sixth-round staining and imaging. The residual signals also appeared in the fifth- and sixth-round images, but we successfully acquired images of the fifth and sixth proteins after unmixing.

## Discussion

In this study, we demonstrated IMPASTO, which enables the cyclic imaging of proteins without signal removal processes. As IMPASTO does not require the inactivation of fluorophores and antibody removal, the staining process can be independently optimized regardless of how challenging signal removal is. Hence, IMPASTO can be used with any fluorophores and antibodies. More importantly, the removal of the signal removal process reduces the epitope loss or tissue distortion across repeated staining and imaging cycles, enabling the volumetric cyclic imaging of thick tissue. In this work, we demonstrated 30-plex imaging in 10 rounds and 12-plex volumetric imaging using IMPASTO. We also demonstrated 30-plex imaging in 5 rounds by using our neural unmixing algorithm to unmix images of spectrally overlapping fluorophores and images from consecutive rounds.

IMPASTO could be combined with existing cyclic imaging techniques to achieve even better multiplexing capabilities or to shorten their signal removal processes. As shown above, the combination of IMPASTO and antibody-stripping-based methods would allow performing more imaging rounds while overcoming the steric hindrance of antibodies. We expect that the steric hindrance of antibodies would be the major factor limiting the maximum number of rounds when imaging dense structures. In such cases, performing an antibody-stripping process once every 10 or 20 rounds would be sufficient to resolve the steric hindrance issues. Even a mild or short antibody-stripping process that can resolve the steric hindrance issues but is not enough for the complete removal of the signals would be sufficient, as the residual signals can be unmixed using IMPASTO. The combination of IMPASTO and chemical-inactivation-based methods could decrease the time needed for the complete removal of fluorescence signals. In addition, IMPASTO would enable the use of extremely photostable fluorophores that could not be used previously in chemical-inactivation-based cyclic imaging techniques.

IMPASTO can be combined with various existing imaging techniques and used for diverse applications. First, IMPASTO can be integrated with signal amplification techniques. Recently, the need to amplify fluorescence signals to detect sparsely expressed proteins has increased^41^. To meet this need, various signal amplification techniques have been developed^24,42,43^. Signals of sparsely expressed proteins could be amplified by one of these techniques without any changes in the IMPASTO process. Second, IMPASTO can be used for imaging cleared or expanded tissues. Tissue-clearing techniques enable the volumetric imaging of thick tissues without thin-sectioning^20,22,38–40,44–48^, while tissue expansion techniques enable the super-resolution imaging of proteins in thick tissue slices^21,31,49–60^. Repeated staining, imaging, and antibody removal have been used in the molecular imaging of cleared^20,21^ or expanded tissues^5,55,61,62^. In such cyclic imaging, the signal removal step could be replaced with IMPASTO’s neural unmixing algorithm. This would greatly reduce the total process time. The combination of IMPASTO with a subset of tissue-clearing techniques, which enable the rapid immunostaining of antibodies, would enable the imaging of proteins over large volumes, such as the whole brain^32,63,64^, without the need for the signal removal process.

## Methods

### Ethical statement

The Korea Advanced Institute of Science and Technology Institutional Animal Care and Use Committee (KAIST-IACUC) approved all the procedures involving animals (approved protocol number KA2020-48).

### Antibodies and chemicals

**Supplementary Data 1–2** list all the antibodies and chemicals used in this study.

### Mouse brain perfusion and slicing

The mice were raised in ventilated cages with a 12-hour light/dark cycle under 20–24°C, 40–60% humidity conditions. Prior to transcardially applying ice-cold 4% PFA in 1× PBS to the mice, we performed isoflurane anesthesia. We extracted the brains and stored them in the same ice-cold 4% PFA solution at 4°C for 2 h before slicing them to the target thickness using a Leica VT1000S vibratome. We stored the sectioned brain slices in 0.1M glycine and 0.01% sodium azide in 1× PBS prior to using them in the experiment.

### PFA-fixed paraffin-embedded brain sample preparation

We dehydrated the PFA-fixed brain samples in an ascending order of 70%, 80%, and 95% ethanol, followed by 100% ethanol twice for 1 h each. We washed the dehydrated brain samples with xylene twice for 1 h each and then immersed the processed samples in paraffin wax and incubated them twice for 1 h each. We embedded the paraffinized brain in the paraffin block mold with melted paraffin wax and cooled it to solidify the residual paraffin wax. We trimmed the paraffin-embedded brain block to 10-um thickness on a microtome and floated the slices in a 45–50°C water bath filled with deionized water. We gently transferred the floated slices to a silanized glass slide with appropriate conditions, as follows: 1) the slices should be fully flattened, without wrinkles that could cause a nonuniform focal plane during imaging, and 2) the slices should be mounted on a slide glass before the tissue could disintegrate. We deparaffinized the slices by washing them twice with xylene for 10 min each. Then, we hydrated the deparaffinized slices with smaller changes in ethanol concentrations to minimize tissue loss during hydration, transferring them to ethanol in descending order as follows: 100% ethanol twice, followed by 95%, 80%, 70%, 60%, and 50% ethanol for 3 min each. We washed the hydrated slices gently with deionized water for another 3 min and then incubated them with heated sodium citrate buffer (pH 8.0) at 80°C for 30 min.

### Conjugation of Fab fragment antibodies with fluorophores

For the Alexa and CF fluorophores, we used a mixture of 90 μL of unconjugated Fab fragment antibody solution, 10 μL of 1M sodium bicarbonate (pH 8.3), and a 9-fold molar excess of succinimidyl ester-fluorophore stock in dimethyl sulfoxide (DMSO). For the ATTO fluorophores, we used a mixture of 90 μL of unconjugated antibody solution, 10 μL of 1M sodium bicarbonate (pH 8.3), and a 3-fold molar excess of succinimidyl ester-fluorophore stock in DMSO. We incubated the solution at room temperature for 1 h in darkness to achieve fluorophore–antibody conjugation and then gently purified the fluorophore-conjugated antibody solution using NAP-5 gel filtration columns. The columns were first washed and equilibrated three times with 3 mL of 1× PBS. We transferred the total volume of fluorophore-conjugated antibody solution (100 μL) into the column, which was then filled with 900 μL of 1× PBS, and we collected the eluates of purified conjugates. We then concentrated the eluates using centrifugal filters with a molecular weight cut-off (MWCO) of 30,000.

### Preparation of preformed antibody complexes

We conducted the primary-Fab complex preformation process based on the original reported procedure^65^. We mixed a solution containing the target primary antibody with 1× PBS and the fluorophore-conjugated Fab fragment antibody in a volume ratio of 10:2:1, and then incubated the mixture for 10 min at room temperature in total darkness. Thereafter, we mixed a five-fold excess of blocking buffer (5% normal rabbit serum, 0.2% Triton X-100, and 1× PBS) and shook it gently for an additional 10 min at room temperature in darkness. All the primary-Fab complexes for each corresponding target protein were combined, diluted in the blocking solution, and used to stain the target sample. We correctly adjusted the concentration of each primary antibody according to the manufacturer’s instructions. We used six fluorophores (two for each laser channel) in this study; CF 488A and ATTO 532 for a 488-nm laser, CF 568 and ATTO 594 for a 557-nm laser, and CF 633 and CF 660R for a 640-nm laser.

### Staining of mouse brain slices with preformed antibody complexes

We incubated the brain slices in a blocking solution (5% normal rabbit serum, 0.2% Triton X-100, and 1× PBS) for 1 h to permeabilize and block the sample. We immunostained the brain slices with a prepared primary-Fab complex mixture overnight in darkness, followed by 30-min washing three times with 0.01% PBST. All these processes were conducted at room temperature.

### Sample distortion issues from the repetitive application of antibody stripping and fluorophore inactivation

We imaged the brain slices before and after the signal removal process under diverse buffer conditions. For antibody stripping, we tested three different stripping buffers based on sodium citrate, a mixture of potassium permanganate and sulfuric acid, and a NewBlot Nitro 5X stripping buffer. For fluorophore inactivation, we tested two different inactivation buffers based on a mixture of hydrogen peroxide and sodium hydroxide and on lithium borohydride (see **Supplementary Data 2** for detailed chemical product information).

### Measurement of intensity decrease trend in fluorophores across staining rounds

We chose six fluorophores to study variations in the intensity decreases of fluorophores across staining rounds. We conjugated Lamin B1 antibody with selected fluorophores (CF 488A, ATTO 532, CF 568, ATTO 594, CF 633, and CF 660R), stained all six fluorophores in individual mouse brain samples, and imaged them across four staining rounds under identical imaging conditions (laser power, exposure time, scan speed, pixel resolution, etc.), as shown in **Supplementary Data 3.**

### Quantitative validation of IMPASTO output images

We based the quantitative validation of IMPASTO on a three-round scale experiment by assigning one of three laser channels for IMPASTO and staining second and third target expressions in the other two laser channels. We quantitatively estimated the IMPASTO unmixing accuracy by comparing IMPASTO output images with reference channel images using Pearson correlation coefficient (r) values (see **Supplementary Data 4**).

### Demonstration of IMPASTO with three standard laser channels

We demonstrated the IMPASTO concept using a single laser channel at an early stage to consider its feasibility for multiplexed cyclic imaging. To extend its multiplexing capability, we simultaneously employed 3 laser channels (except the 405-nm laser channel) as IMPASTO channels.

### 3D IMPASTO imaging

For 3D IMPASTO imaging, we performed 20 μm-depth z-stack imaging with a 1.0-μm step size, meaning that we acquired 21 images per laser channel (84 images in total). Since each z-slice exhibited a slightly different degree of deformation, we had to acquire and apply an image registration vector map for each z-slice.

### Spectro-temporal unmixing via IMPASTO

IMPASTO is a method of selectively unmixing the target expression between the “A” and “A + B” images. Therefore, the IMPASTO output would be an image of “A” itself and an image of “B.” Specifically, the general hypothesis is that the unmixing matrix for IMPASTO should be in the form 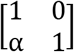, where α is less than 1. However, in spectral unmixing applications, it is necessary to unmix spectrally overlapped signals, such as images of “A + b” and “a + B,” where the capital letter is the dominant case in the corresponding detection range and vice versa. Such a statement implies that the unmixing matrix in spectral unmixing should be in the form 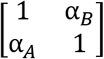, where α*_A_* and α*_B_* are both less than 1.

### IMPASTO conjunction with the conventional antibody stripping method

After a three-round scale IMPASTO experiment, we performed antibody stripping with a NewBlot Nitro 5x stripping buffer for 10 min per round. Unlike thick brain slice samples, paraffin-embedded samples were comparatively more stable regarding sample distortion from antibody stripping. However, we observed that repetitive antibody stripping can cause sample distortion, as well as detachment of paraffin-embedded slices from the slide glass (see **Supplementary Figure 13** and **Supplementary Video 4**). Therefore, it made sense to periodically apply mild antibody stripping in the IMPASTO experiment in several tens of staining rounds for much further staining rounds without steric hindrance issue.

### Fluorescent imaging and visualization

In this study, we performed most of the experiments, including the quantitative/qualitative analyses and 30-plex fluorescent imaging, using a Nikon C2 Plus point-scanning confocal microscope equipped with four excitation lasers (405, 488, 557, and 640 nm) and a spectral detector. We acquired all the imaging data using a water immersion 40x objective lens with 1.15 NA. We conducted antibody screening, 3D IMPASTO imaging, and IMPASTO with antibody stripping experiments using a spinning-disk confocal microscope from Andor DragonFly equipped with four excitation lasers (405, 488, 561, and 640 nm) with a normal filter-based detection system. We acquired all the imaging data from Andor DragonFly using a water-immersion 40x objective lens with 1.10 NA. The brightness of the images included in the figures, Supplementary Figures, and Supplementary Videos was adjusted using the “Auto Adjust” function of ImarisViewer software (ImarisViewer 9.9.1) to improve visibility. For the images shown in **Figure 6j** and **Supplementary Figure 17**, the level of autofluorescence and non-specific binding was manually determined for each protein and subtracted from the whole images for better visibility. For images for quantitative analysis, such as the measurements of the unmixing accuracy of IMPASTO, no thresholding was applied to the unmixed images or ground-truth images.

### Image registration and intensity normalization

We processed the image registration using the ImageJ plugin “bUnwarpJ.” The DAPI image in the first staining round was the reference fiducial marker image, whereas the DAPI images in the other staining rounds were the images to be registered. For image registration in each staining round, we used the DAPI images from both the first round and the target *N*^th^ round to extract the corresponding registration vector maps. We applied these registration vector maps to other IMPASTO channels to finely match the coordinates of the staining rounds. In 3D IMPASTO imaging, the absolute intensity gradually decreased along with an increase in the imaging depth of the sample. Thus, the intensity scale of the IMPASTO output image for each z-slice was normalized using customized MATLAB code, assigning the first slice (the brightest slice) as the reference intensity scale.

### Neural unmixing for blind signal separation

In neural unmixing, the input image is fed into the unmixing layer, which is a linear unmixing matrix with bias subtraction, to calculate the unmixed image. The pixels in the unmixed image are sampled from the joint distribution and the product of the marginal distributions. Both samples were then passed to the independence evaluation network, which measures the independence among the channels.

To ensure the permutation invariance of the independence evaluation network (i.e., the order of sampled pixels in each set should not affect the independence assessment), the network is designed to have a permutation invariance property based on the following network architecture:

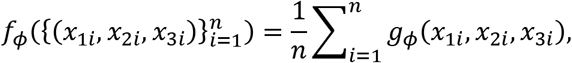

where *g_ϕ_*(·) is a multilayer perceptron parameterized by *ϕ* that takes only one sample (*x*_1_, *x*_2_, *x*_3_) as the input. The independence evaluation network is trained by minimizing the binary cross-entropy loss 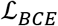 and the unmixing layer is trained by minimizing the composite loss function 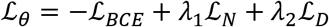 where 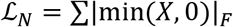 is the negativity loss, 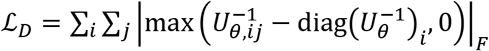 is the diagonal loss that prevents the off-diagonal elements of the mixing matrix from becoming greater than the diagonal elements, *U_θ_* denotes the parameterized unmixing matrix, |·|_*F*_ denotes the Frobenius norm, and *λ*_1_, *λ*_2_ are the regularization parameters. We increased the parameters gradually over the iterations. Using these loss functions, we trained the unmixing layer and the independence evaluation network in an adversarial manner: the classifier network was trained to classify independent and dependent samples, and the unmixing layer was updated to make them indistinguishable while retaining the non-negativity of the unmixed image.

Each multi-layer perceptron *g_ϕ_*(·) has two hidden layers that employ LeakyReLU^66^ with a slope of 0.2 as the activation function. We implemented the adversarial training of the unmixing layer and the independence evaluation network by inserting a gradient reversal layer^67^ between them. The gradient reversal layer multiplied the gradient by –γ during the backpropagation, where the minus sign flipped the sign of the gradient and the value γ = 5 determined the relative learning ratios for the unmixing layer and the independence evaluation network. For training the network, we used an Adam optimizer^68^ with a learning rate of 0.0001 and exponential decay rates of 0.9 for the first moment and 0.999 for the second moment. We implemented the network using PyTorch^69^ and performed all the computations on a workstation with two Intel^®^ Xeon^®^ Silver 4214R CPUs, a 256GiB RAM, and an NVIDIA GeForce RTX 2080 Ti GPU.

For inter-round channel unmixing of two images, when the first image contained the fluorescent signals that accumulated from the first round to the previous round and the second image contained the linearly scaled version of the first image and the fluorescent signal from the current round, we set the unmixing matrix as a 2 × 2 lower triangular matrix (i.e., 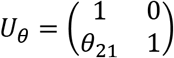) to accelerate the training procedure.

### Neural unmixing of an arbitrary number of channels

Through the simple generalization of the three-channel unmixing algorithm, neural unmixing can separate input images with an arbitrary number of channels. For *N*-channel unmixing, the first set ***x*** is sampled from the joint distribution 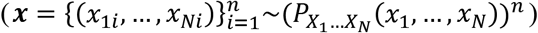, and the second set ***x**′* is sampled from the product of the marginal distributions 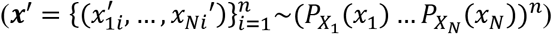, where 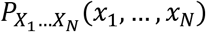 is the joint probability distribution of *X*_1_, …, *X_N_*, and 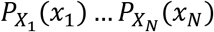 is the product of the marginal distributions. As with the three-channel unmixing case, we trained the classifier network to classify independent and dependent samples by minimizing 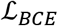, and we updated the unmixing layer to make them indistinguishable by maximizing 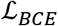. While this formulation was mathematically appropriate for extension to unmixing an arbitrary number of channels, such training in practice can suffer from numerical stability issues due to the “curse of dimensionality.” Thus, we took a divide-and-conquer approach to alleviate this issue.

From the unmixing-layer perspective, maximizing the binary cross-entropy loss is mathematically equivalent to minimizing the Jensen–Shannon divergence between the joint distribution and the product of marginal distributions^70^ :

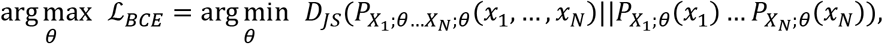

where *θ* are the parameters of the unmixing layer, and the 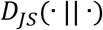 denotes the Jensen–Shannon divergence between two input probability distributions. Applying the divide-and-conquer principle to this optimization problem, we constructed the loss function 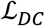 as an upper bound of the square root of the Jensen–Shannon divergence between the joint distribution and the product of marginal distributions, the global minimum of which coincided with that of the original loss function:

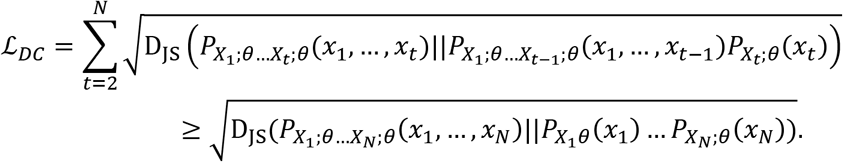

Estimating this loss function required *N* – 1 independence evaluation networks, each of which attempted to distinguish the samples from 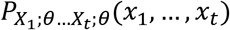 and 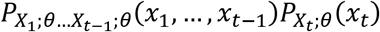, where *t* is an integer between 2 and *N*. In this setting, each independence evaluation network had to distinguish only two distributions that could differ along only one dimension, facilitating the convergence of the unmixing layer.

### Self-supervised image denoising network

As a preprocessing step before unmixing, we performed self-supervised image denoising using convolutional neural network architecture with the blind-spot property proposed in ^71^. The network consisted of three components (see **Supplementary Figure 22a**): (1) a 3 × 3 convolutional path, (2) a blind-spot convolutional path, and (3) a 1 × 1 convolutional path. In the 3 × 3 convolutional path, the input was passed through 10 sequential convolutional layers with a 3 × 3 kernel size. The feature maps generated in the 3 × 3 convolutional path were passed to the blind-spot convolutional path, the receptive field of which was set as zero at its center. The outputs from the blind-spot convolutional layers were concatenated and passed through three 1 × 1 convolutional layers to predict the denoised image while maintaining the blind-spot property. The number of channels was set as 32 for the 3 × 3 convolutional path and 64 for the blind-spot convolutional path. In the 1 × 1 convolutional path, the numbers of channels were set as 64, 16, and 2 for each layer. The receptive field of the network is visualized in **Supplementary Figure 22b**, which shows the blind-spot property. We trained the network to minimize pixel-wise loss, defined as a linear summation of L_1_ and L_2_ loss between the input and the output. To train the network, we used an Adam optimizer^68^ with a learning rate of 0.0003. We performed all computations on a workstation with two Intel^®^ Xeon^®^ Silver 4214R CPUs, a 256GiB RAM, and an NVIDIA GeForce RTX 2080 Ti GPU.

## Acknowledgments

The Korean National Research Foundation (NRF) supported this work with a grant funded by the Korean government (MSIT) (NRF-2021R1C1C1006642, NRF-2020R1C1C1009869), and the Bio & Medical Technology Development Program of the NRF supported it with a grant funded by MSIT (NRF-2021M3A9I4026318).

## Author Contributions

J.-B. C., Y.-G. Y., H. K., and S. B. conceived the main idea. H. K. and J. S. optimized the experimental protocol and performed immunostaining and imaging. H. K. conducted image registration and quantitative analyses. H. N. performed the antibody screening. S. B., J. C., S. H., E. Y., and E. K. wrote all the unmixing and image-processing programs. All authors drafted and contributed to the editing of the paper. J.-B. C. and Y.-G. Y. supervised this work.

## Data Availability

The data to support the findings of this study are available from the corresponding author upon request.

