## Supplementary Information for "IMPASTO: Multiplexed cyclic imaging without signal removal *via* self-supervised neural unmixing"

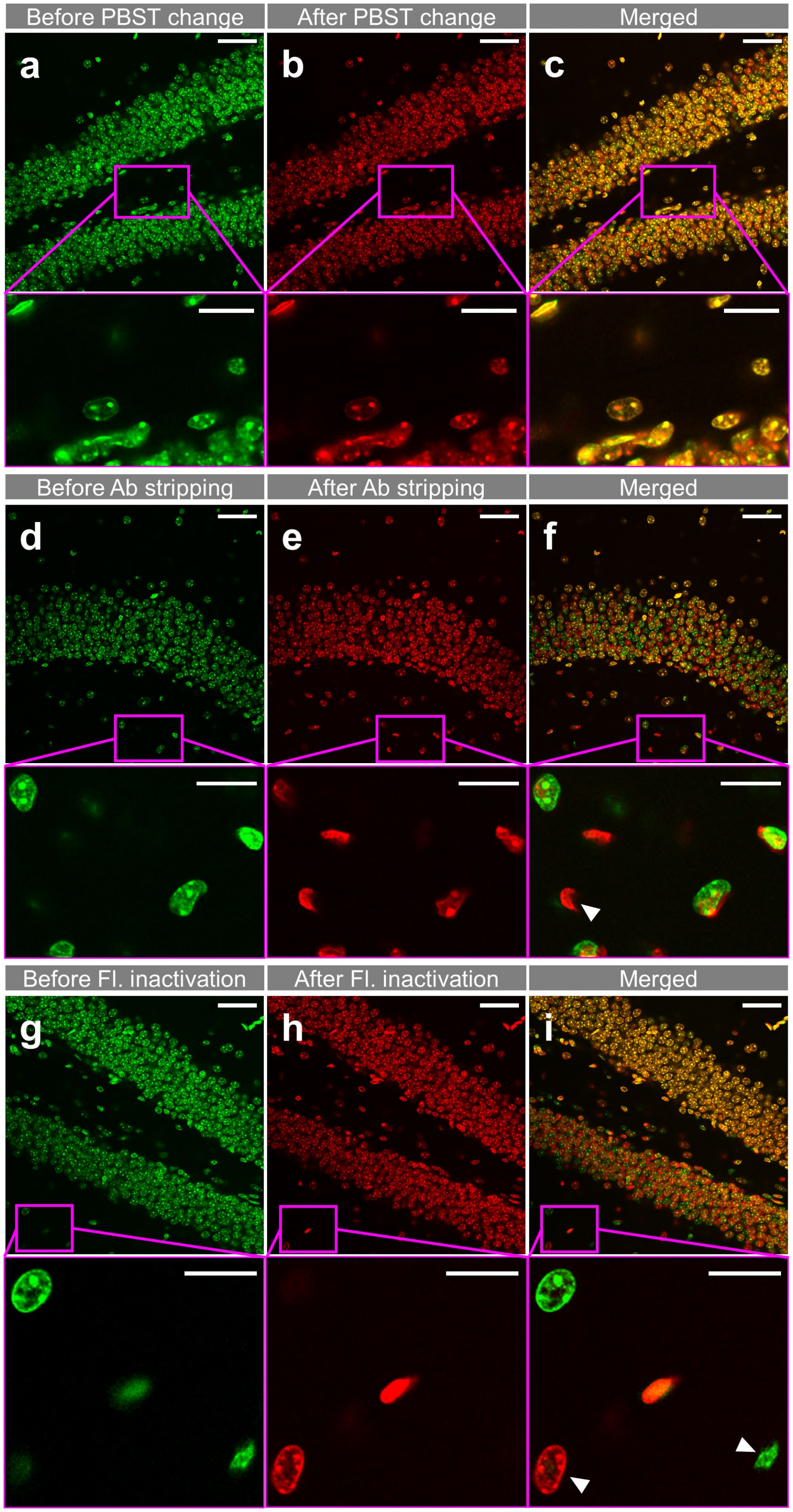
Before PBST change
After PBST change
Merged
a
b
c
Before Ab stripping
After Ab stripping
Merged
d
e
f
Before Fl. inactivation
After Fl. inactivation
Merged
g
h
i

**Supplementary Figure 1. Sample distortion after 10 rounds of buffer exchange.** Images of 50-μm-thick mouse brain slices processed with three different buffer exchange procedures for 10 rounds. (**a–c**) In each round, a brain slice was incubated in a wash buffer (1× PBS, 0.2% Triton X-100) for 10 min and then washed with the same buffer for 3 min. This process was repeated for 10 rounds. (**a**) Before the buffer exchange. (**b**) After the 10 rounds of buffer exchange. (**c**) Merged view of the images shown in **a** and **b**. (**d–f**) In each round, a brain slice was incubated in a fluorophore-inactivation reagent[1] for 10 min and then washed with a wash buffer for 3 min. This process was repeated for 10 rounds. (**d**) Before the buffer exchange. (**e**) After the 10 rounds of buffer exchange. (**f**) Merged view of the images shown in **d** and **e**. (**g–i**) In each round, a brain slice was incubated in NewBlot Nitro 5X stripping buffer for 10 min and then washed with a wash buffer for 3 min. This process was repeated for 10 rounds. (**g**) Before the buffer exchange. (**h**) After the 10 rounds of buffer exchange. (**i**) Merged view of the images shown in **g** and **h**. Arrowheads shown in the magnified views of **f** and **i** show mismatches between the before and after buffer exchange images. Such mismatches are the results of the axial distortion of the brain slices. Scale bar = 50 μm in (**a-i**) and 20 μm in the magnified views.

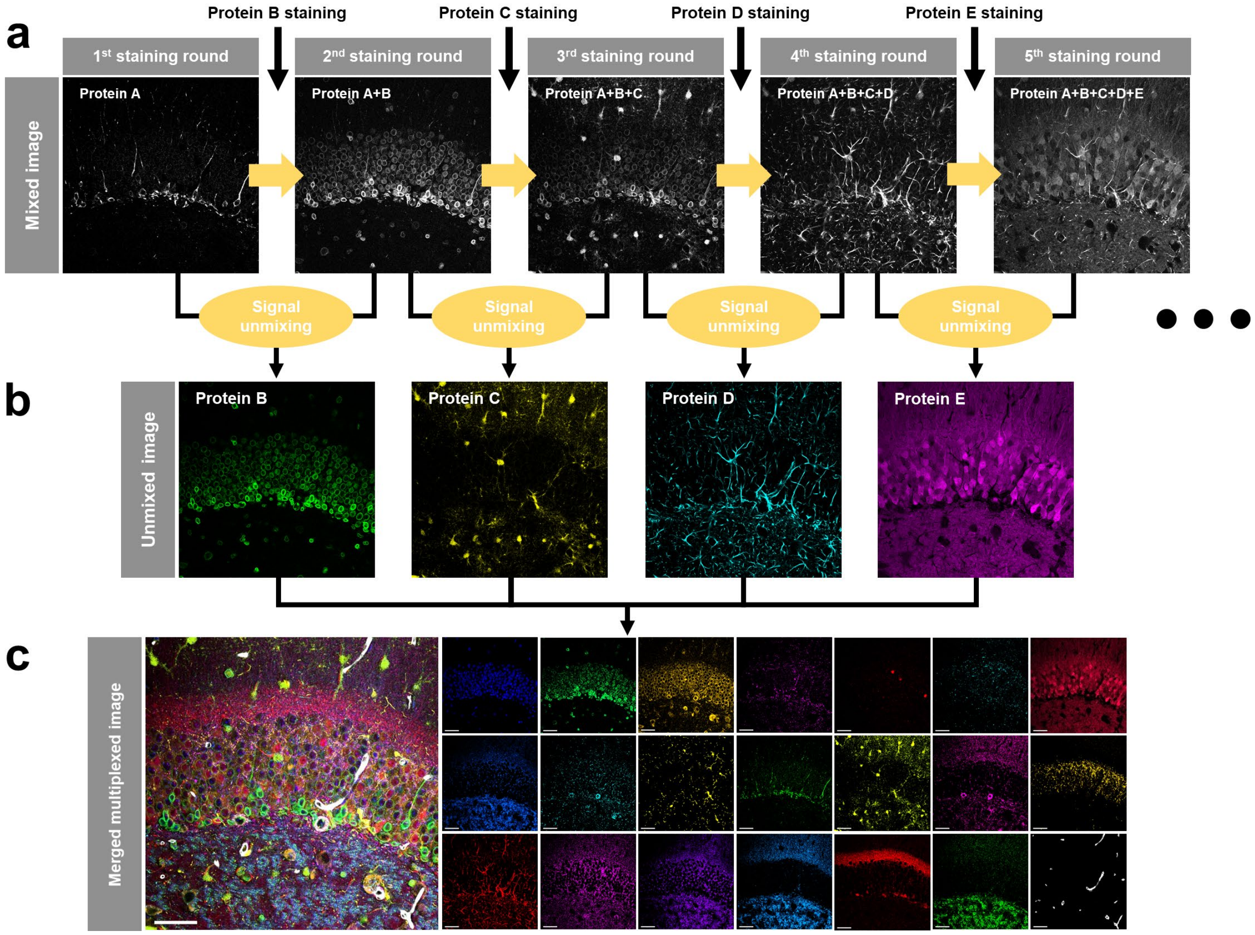


**Supplementary Figure 2. A process of acquiring 5-plex images using IMPASTO.** (**a**) Staining process. A specimen is iteratively stained and imaged without a signal removal process. (**b**) Images of single proteins are retrieved by the unmixing images of two consecutive rounds. (**c**) An exemplary result of multiplexed cyclic imaging *via* IMPASTO. Scale bar = 50 μm.

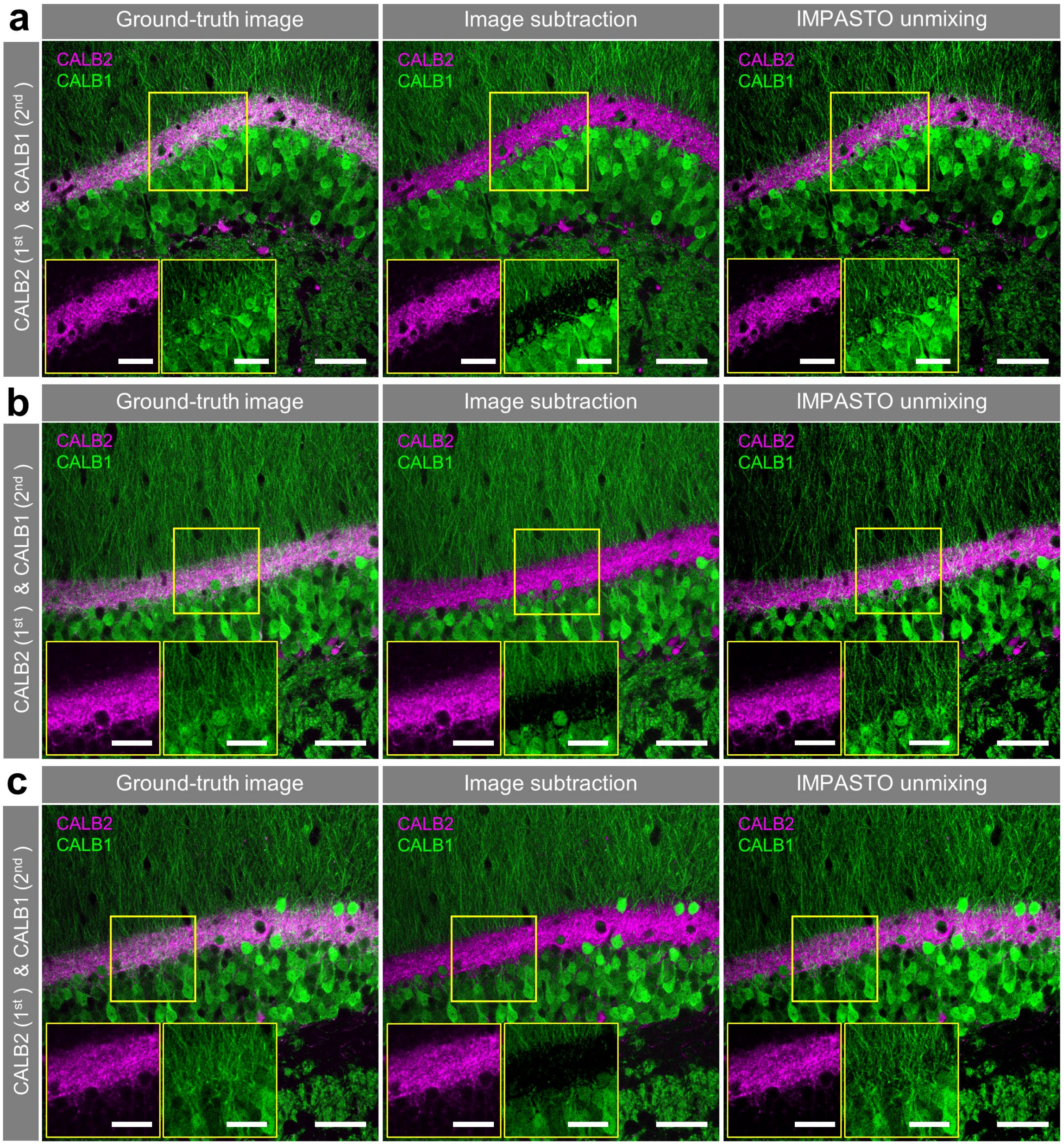


**Supplementary Figure 3. Three more examples of the unmixing of images acquired from two consecutive rounds using measured α and our neural unmixing code.** 50-μm-thick mouse brain slice was stained against CALB2 in the first round and against CALB1 in the second round, using the same fluorophore. Simultaneously, these two proteins were stained with spectrally distinctive fluorophores, and their images were used as ground-truth images. (**a**–**c**) Left panel: ground-truth images of the two proteins. Center panel: results of the unmixing of the first- and second-round images using α shown in **Figure 1d**. Note that the CALB2 signal was over-subtracted from the CALB1 signal. Right panel: results of the unmixing of the first- and second-round images using our neural unmixing algorithm. The results of IMPASTO unmixing (right panels) matched the ground truth images (left panels). Scale bar = 50 μm in (**a**-**c**) and 20 μm in the magnified views.

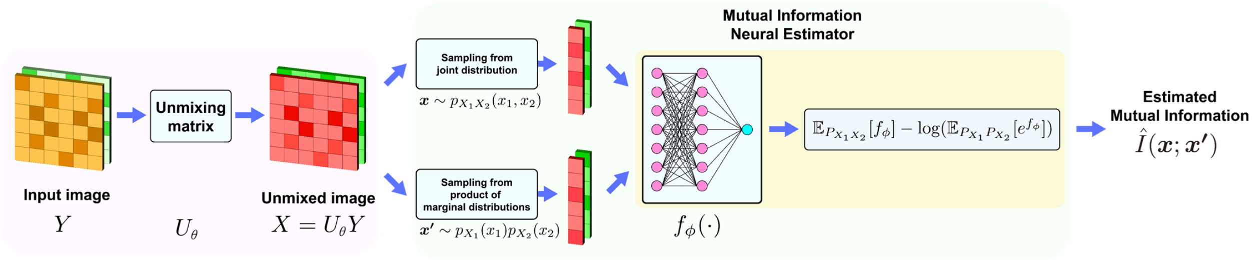


**Supplementary Figure 4. IMPASTO using mutual information neural estimator (MINE) as the independence evaluation network.** Similar to the IMPASTO that employs a classifier network as the independence evaluation network, the unmixing matrix and the MINE are trained in an adversarial manner; the mixing matrix is trained to minimize the estimated mutual information whereas the MINE is trained to maximize it.

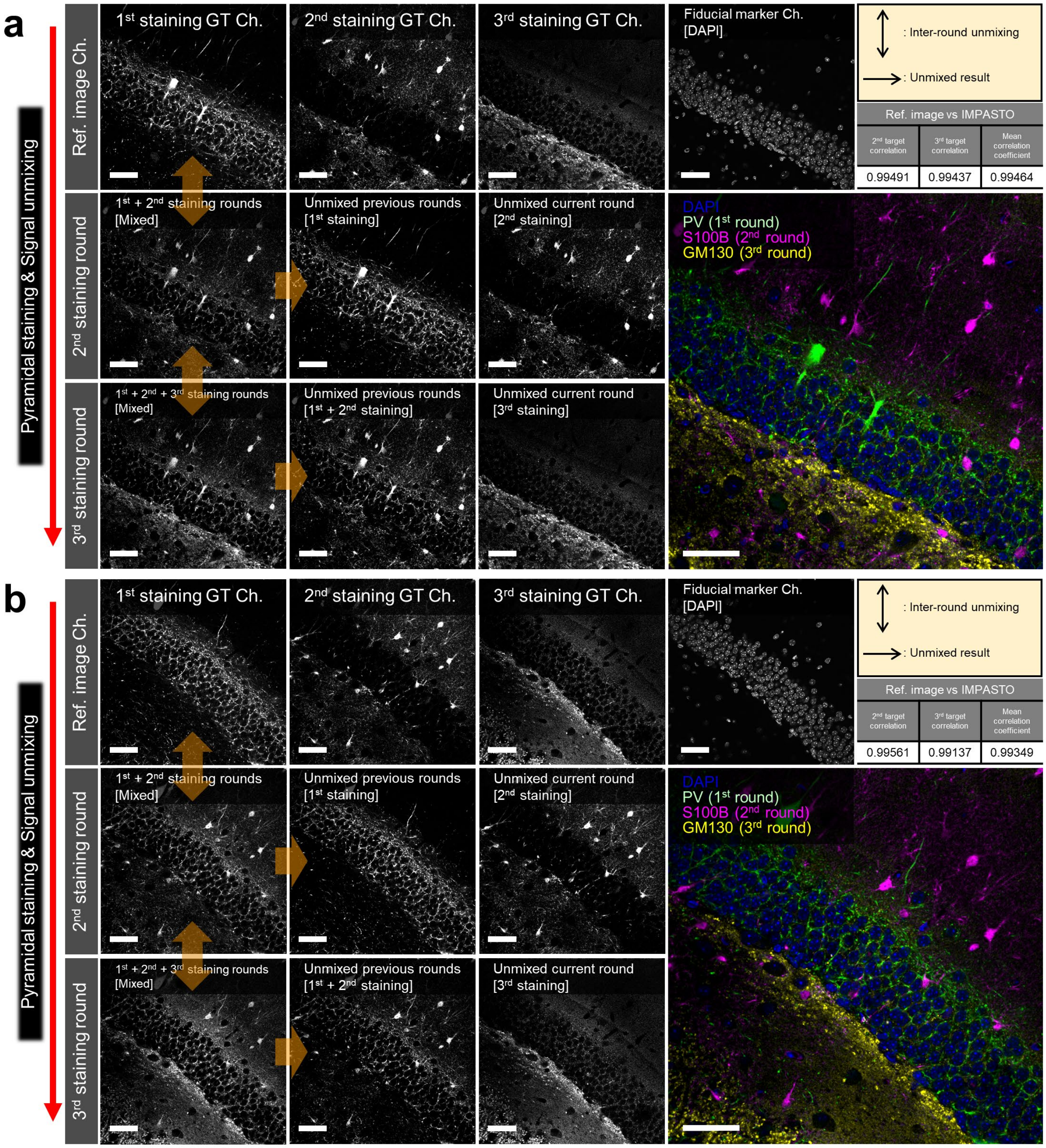


**Supplementary Figure 5. Quantitative analysis of the unmixing accuracy of IMPASTO.** (**a**–**b**) Two validation studies performed on different brain samples. Target proteins were PV ($1_{st}$ round), S100B ($2_{nd}$ round), and GM130 ($3_{rd}$ round). Unmixing accuracy was estimated by measuring Pearson correlation coefficients between the unmixed and ground-truth images. Pearson correlation coefficients were greater than 0.99 for all proteins in both experiments. Scale bar = 50 μm.

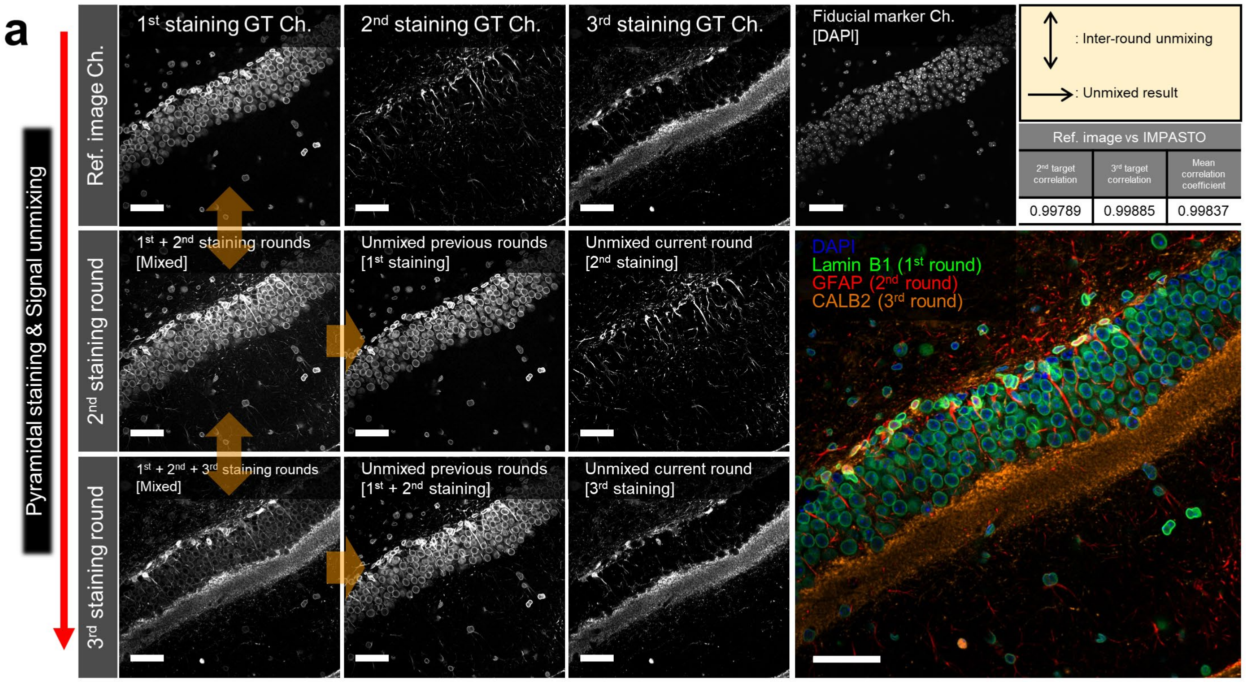


**Supplementary Figure 6. Quantitative analysis of the unmixing accuracy of IMPASTO with different antibody panels.** Target proteins were lamin B1 ($1_{st}$ round), GFAP ($2_{nd}$ round), and CALB2 ($3_{rd}$ round). Unmixing accuracy was estimated by measuring Pearson correlation coefficients between the unmixed and ground-truth images. Pearson correlation coefficients were greater than 0.99 for all proteins. Scale bar = 50 µm.

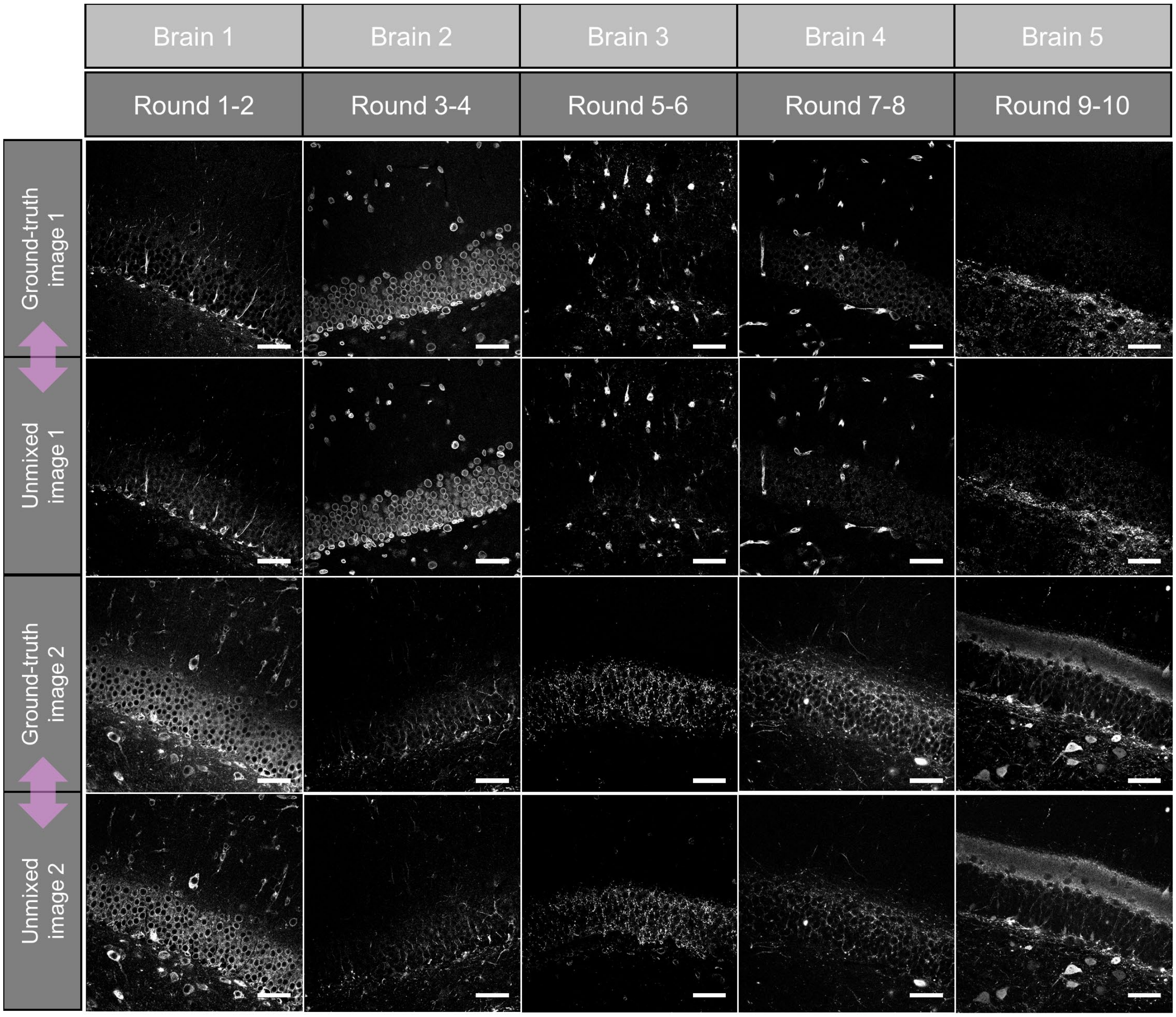


**Supplementary Figure 7. Quantitative analysis of the unmixing accuracy of IMPASTO over 10 rounds.** 10-round cyclic staining and imaging was performed on five brain slices. Each brain slice was simultaneously stained with two of ten antibodies and their images were used as ground-truth images. The unmixing results were compared with respective ground-truth images. For example, the unmixed images of the fifth and sixth proteins were acquired from brain 3 and compared with their ground-truth images acquired from the same brain. Note that in all five brain slices, the unmixed images match their ground-truth images. Scale bar = 50 μm.

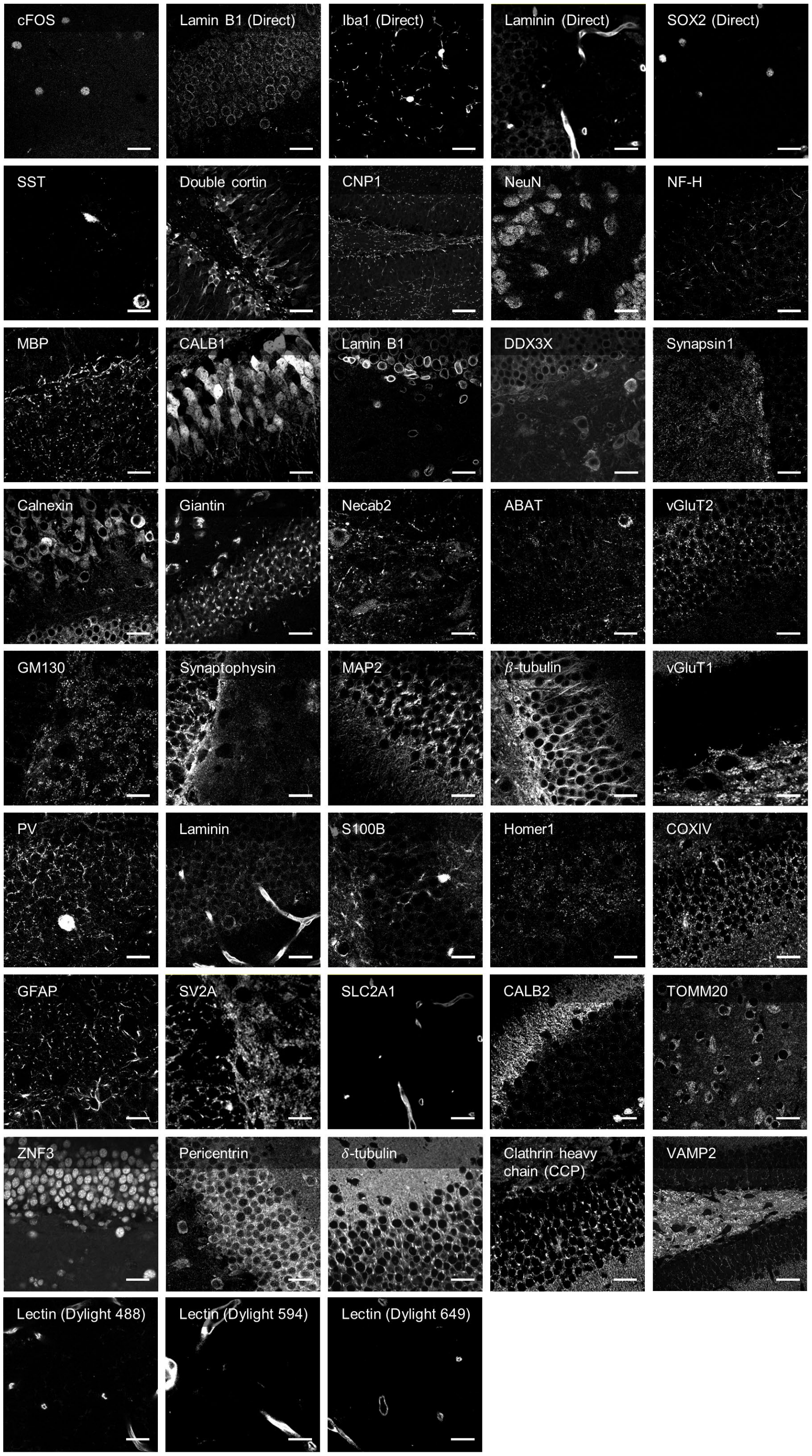


**Supplementary Figure 8. Images of singly-stained mouse brain slices.** The protein expression patterns from singly-stained brain slices were qualitatively compared with the 30-plex image shown in **Figure 4**. Scale bar = 50 μm.

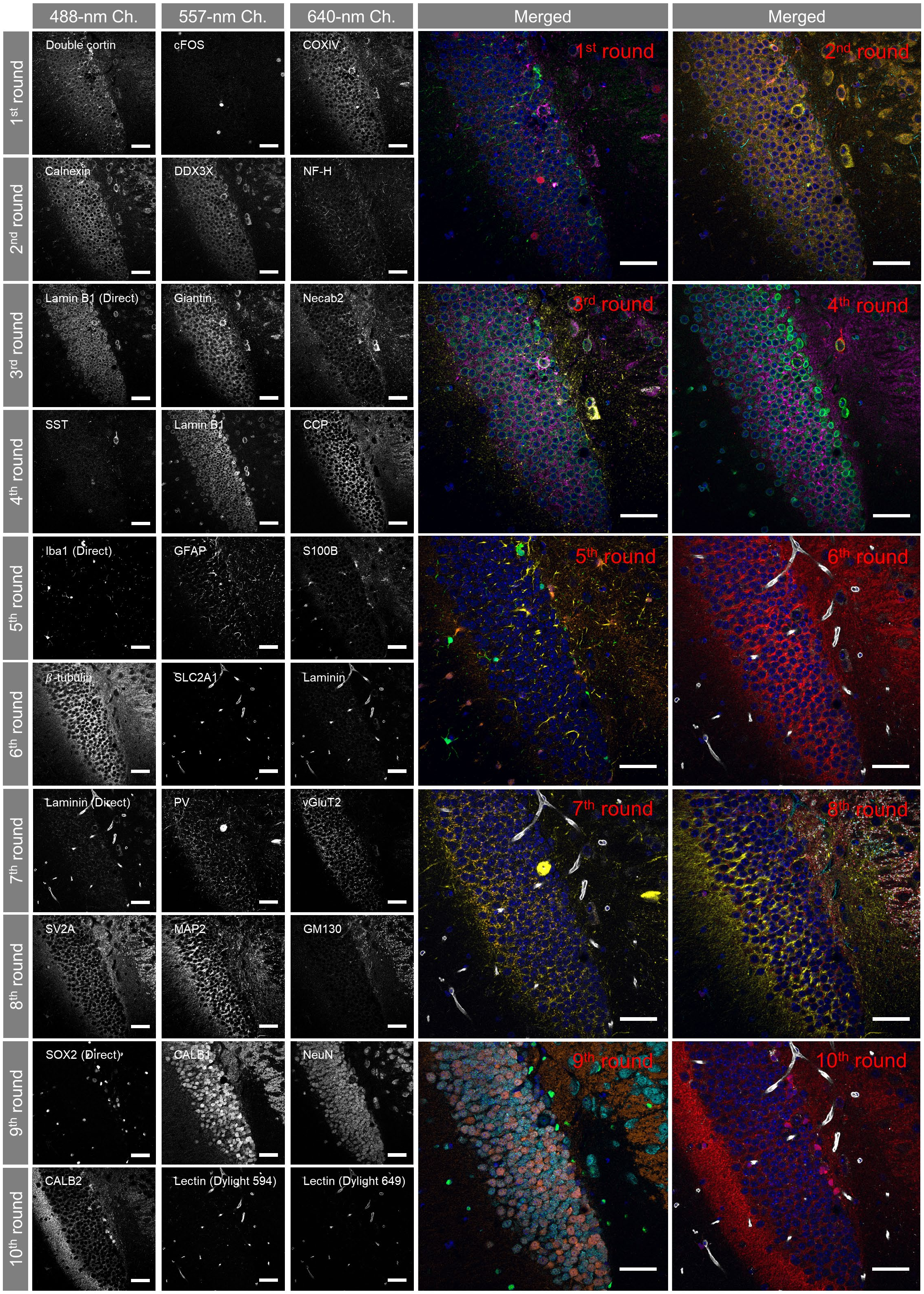


**Supplementary Figure 9. Single-protein images of the 30-plex result shown in Figure 4.** Scale bar = 50 μm.

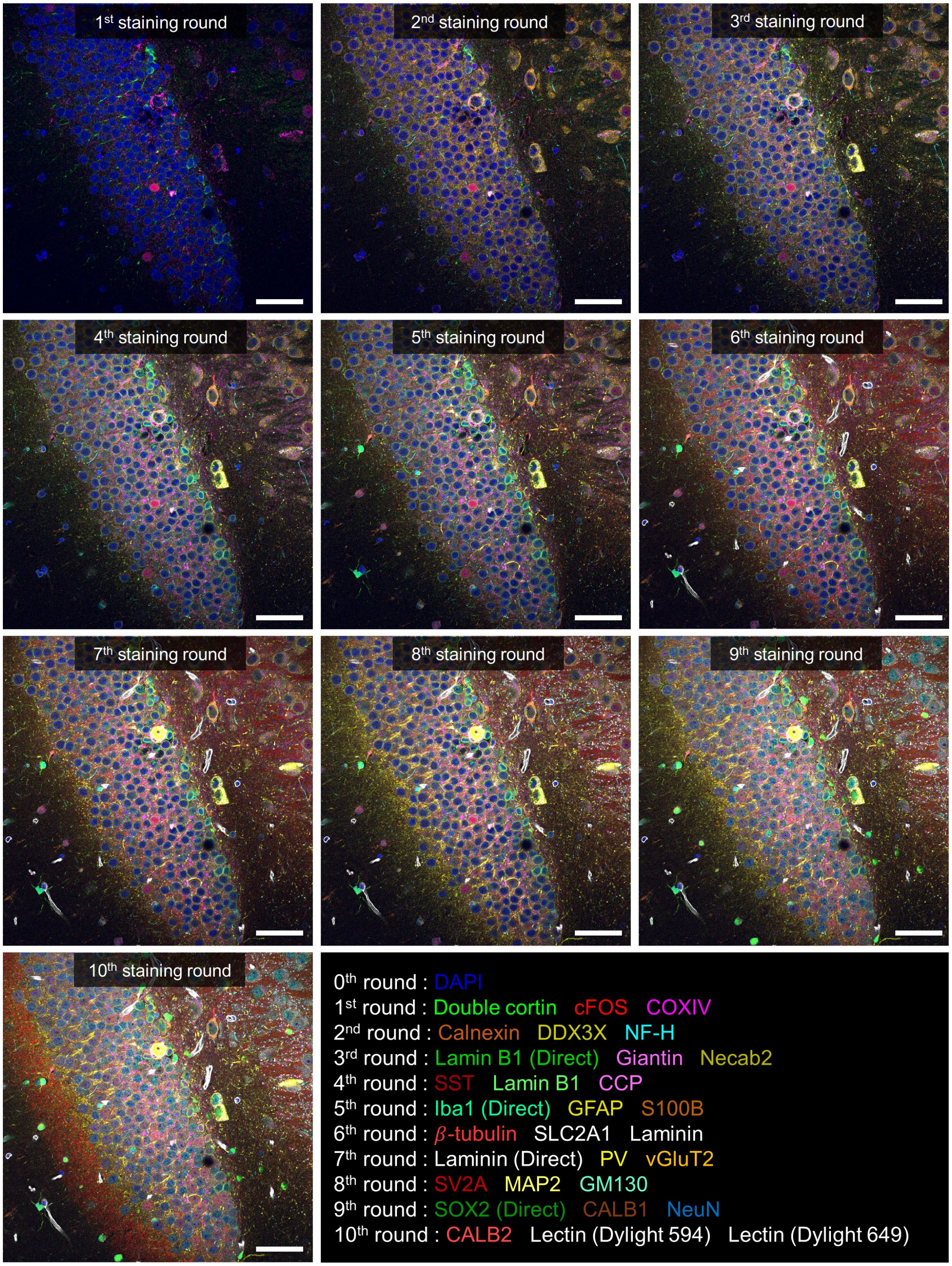


**Supplementary Figure 10. Cumulative visualization of the 30-plex result shown in Figure 4.** Scale bar = 50 μm.

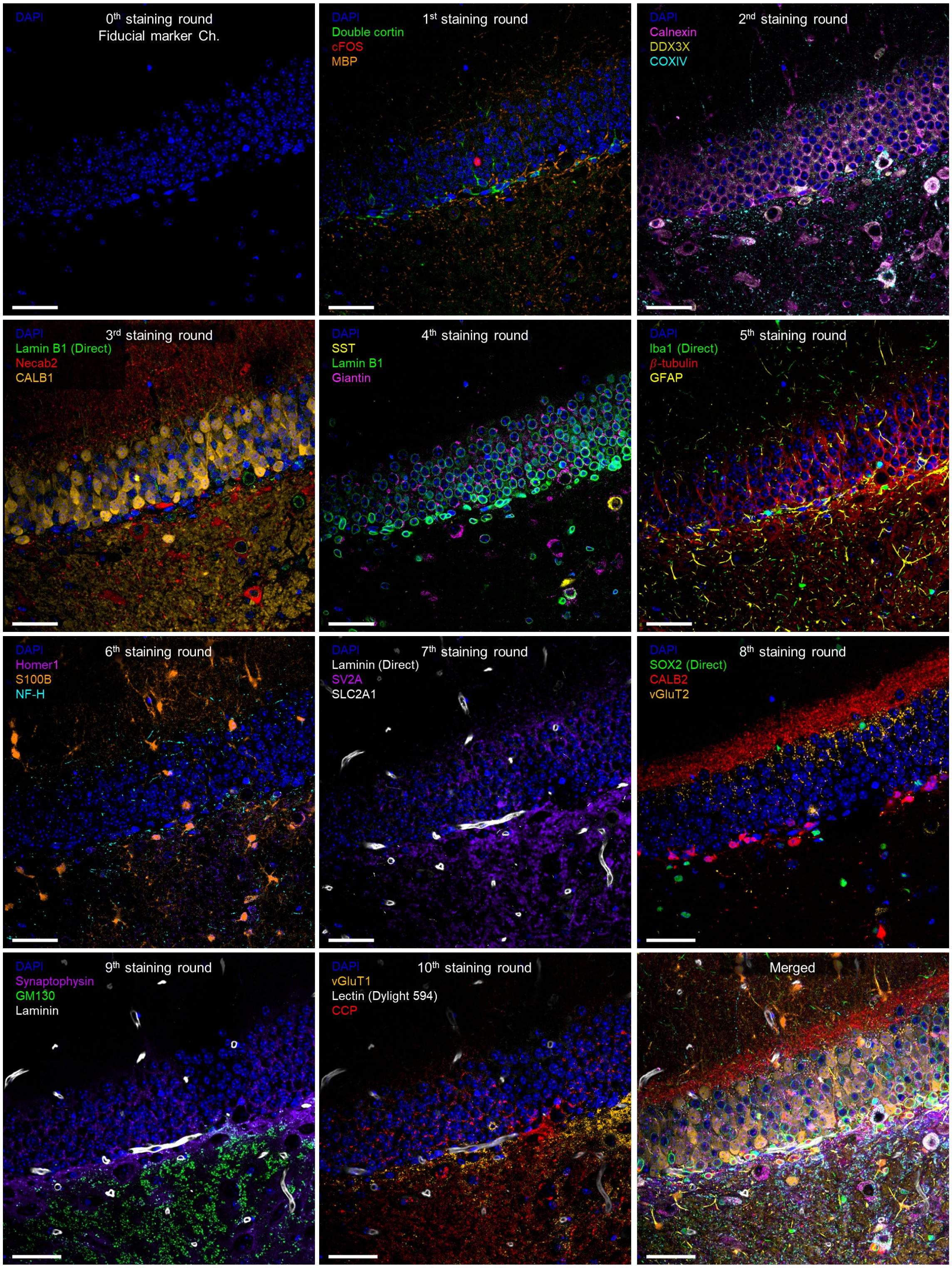


**Supplementary Figure 11. An additional demonstration of highly multiplexed cyclic imaging with different antibody panels.** 30-plex imaging was performed in 10 rounds. DAPI staining was used as a fiducial marker. Scale bar = 50 μm.

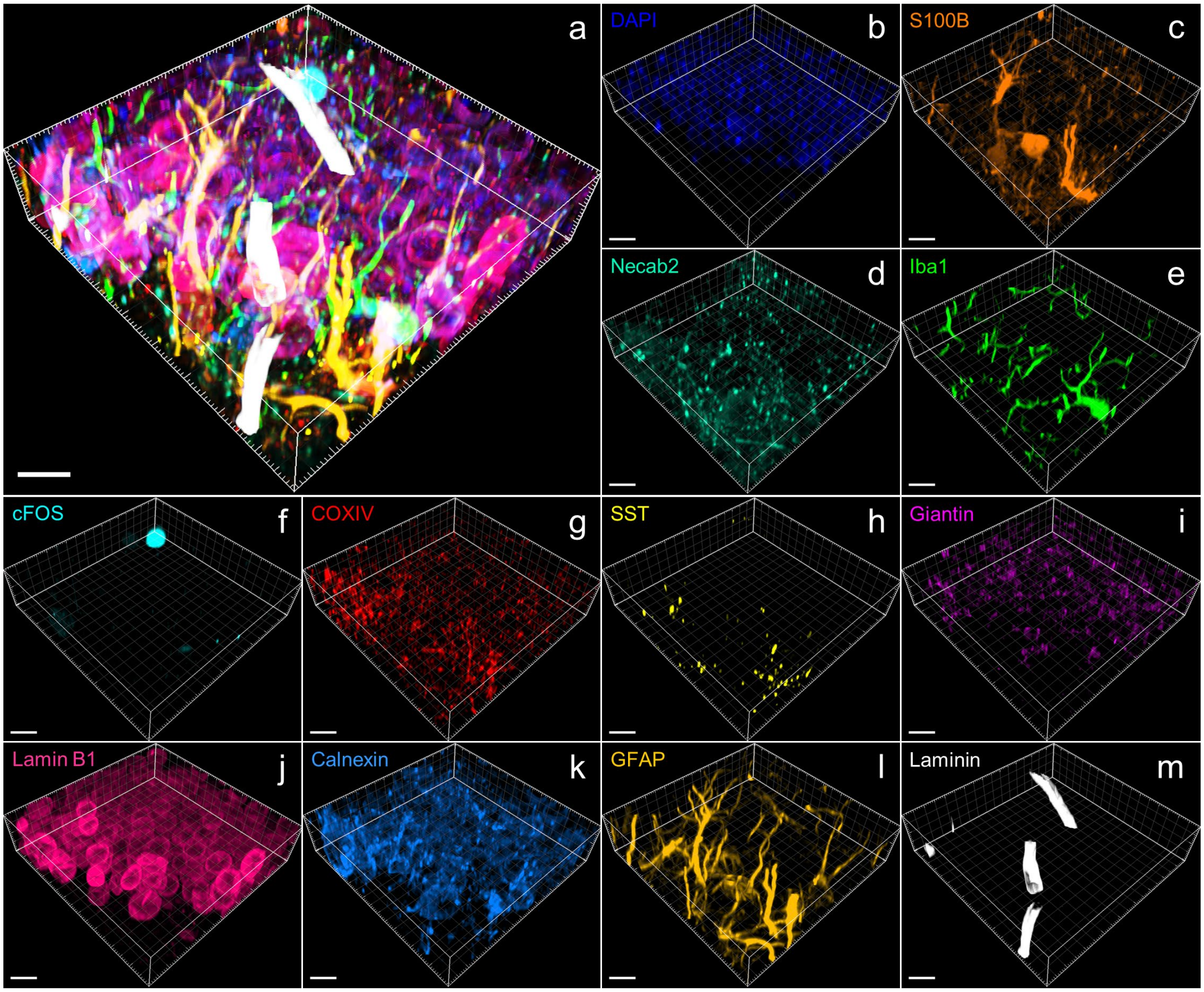


**Supplementary Figure 12. Magnified view of the 12-plex 3D result shown in Figure 5.** (**a**) Merged image of the 12 proteins. (**b-m**) Single-protein images. Scale bar = 10 μm.

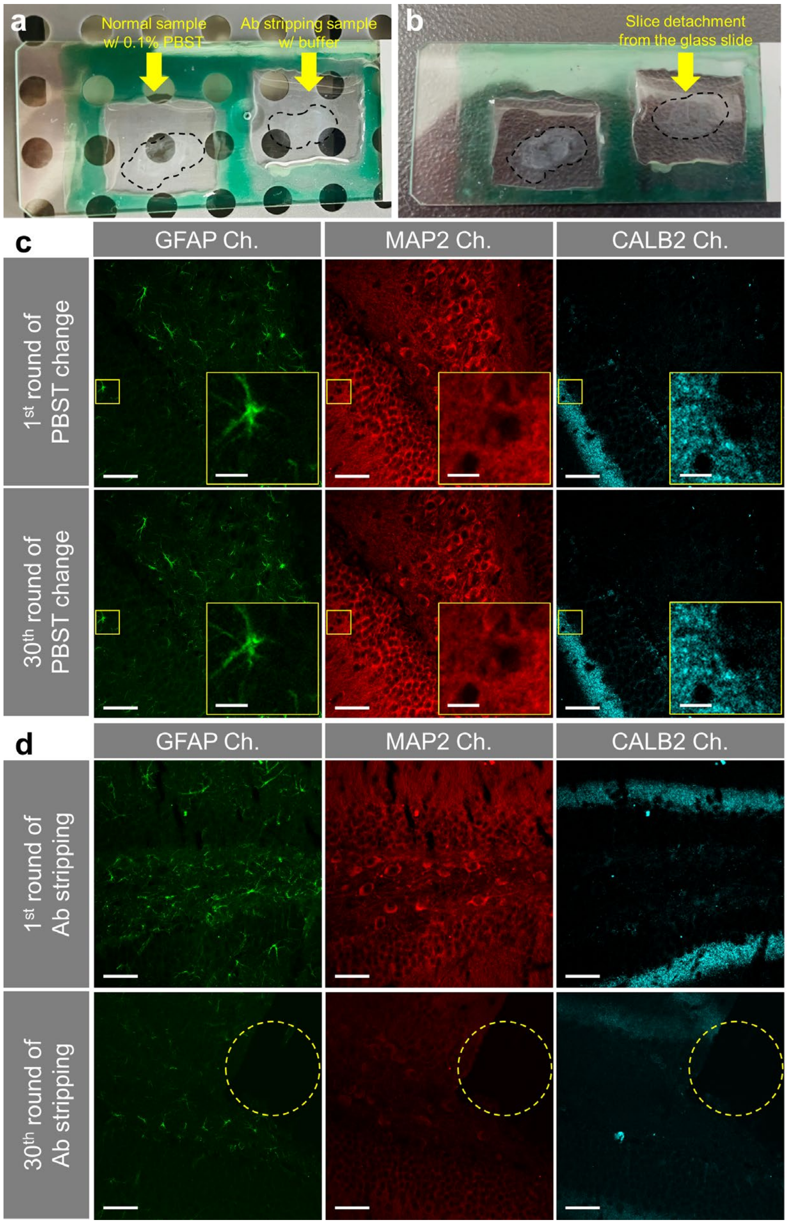


**Supplementary Figure 13. Formalin-fixed paraffin-embedded (FFPE) mouse brain slices after 30 rounds of an antibody-stripping process.** (**a**–**b**) Two FFPE mouse brain slices were stained with antibodies against GFAP, MAP2, and CALB2. Then, these slices were processed with two different treatments for 30 rounds. Each round consisted of the incubation of the slices either in 0.1% PBST (left slice) or NewBlot Nitro 5X stripping buffer (right slice) for 10 min, followed by washing both slices with 0.1% PBST three times, 3 min each. This process was repeated for 30 rounds. (**a**) Before the treatments. (**b**) After 13 rounds of treatments. Note that the right slice was detached from the glass slide. (**c**) Images of the three proteins in the left brain slice before and after the 30 rounds of treatments. Note that no tissue distortion is observed, and the antibody signals are well conserved. (**d**) Images of the three proteins in the right brain slice before and after the 30 rounds of treatments. For this slice, antibodies were applied again after the 30 rounds of treatments. Note that a portion of the tissue is missing (dotted circle) and the signal intensity has decreased. Scale bar = 50 μm in (**c**–**d**) and 10 μm in the magnified views.

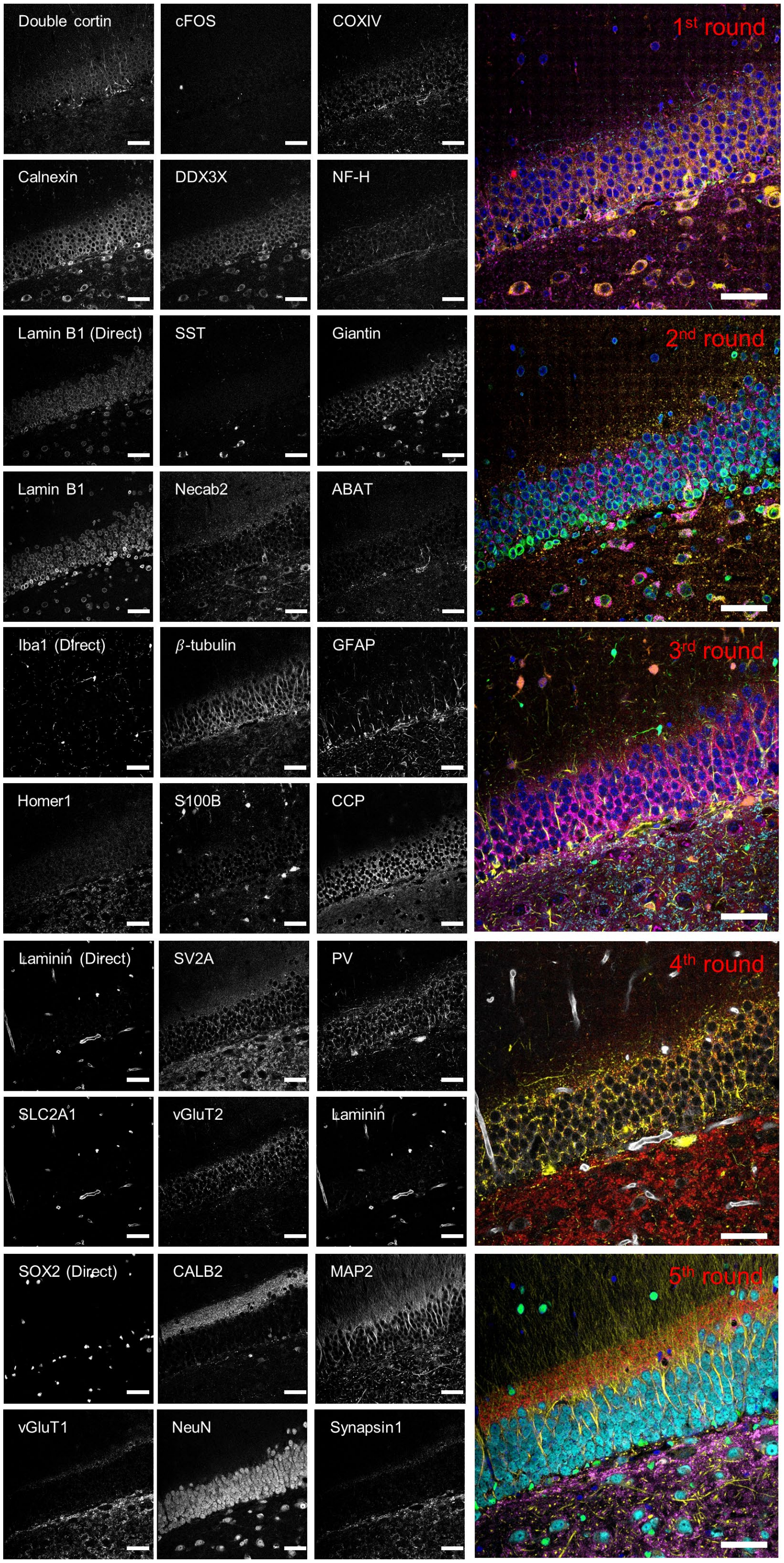

Double cortin
cFOS
COXIV
1st round
Calnexin
DDX3X
NF-H
Lamin B1 (Direct)
SST
Giantin
2nd round
Lamin B1
Necab2
ABAT
Iba1 (Direct)
β-tubulin
GFAP
3rd round
Homer1
S100B
CCP
Laminin (Direct)
SV2A
PV
4th round
SLC2A1
vGluT2
Laminin
SOX2 (Direct)
CALB2
MAP2
5th round
vGluT1
NeuN
Synapsin1

**Supplementary Figure 14. Single-protein images of the spectro-temporal unmixing result shown in Figure 6.** 30-plex imaging was performed in 5 rounds. Two spectrally overlapping fluorophores were used for each of the 488-, 557-, and 640-nm excitation lasers, and their signals were unmixed using the neural unmixing algorithm. Scale bar = 50 μm.

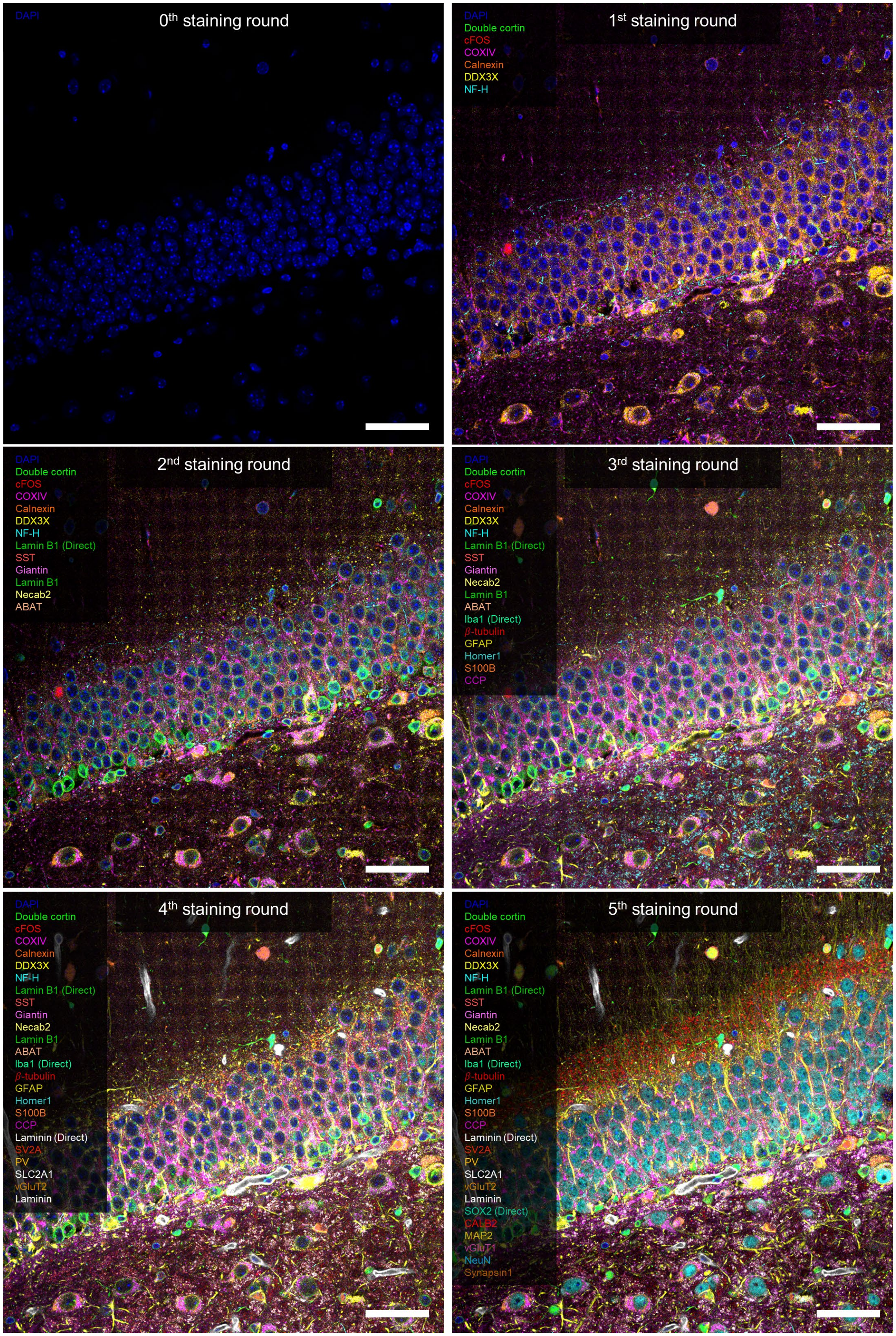
0th staining round
DAPI
1st staining round
DAPI
Double cortin
cFOS
COXIV
Calnexin
DDX3X
NF-H
2nd staining round
DAPI
Double cortin
cFOS
COXIV
Calnexin
DDX3X
NF-H
Lamin B1 (Direct)
SST
Giantin
Lamin B1
Necab2
ABAT
3rd staining round
DAPI
Double cortin
cFOS
COXIV
Calnexin
DDX3X
NF-H
Lamin B1 (Direct)
SST
Giantin
Necab2
Lamin B1
ABAT
Iba1 (Direct)
β-tubulin
GFAP
Homer1
S100B
CCP
4th staining round
DAPI
Double cortin
cFOS
COXIV
Calnexin
DDX3X
NF-H
Lamin B1 (Direct)
SST
Giantin
Necab2
Lamin B1
ABAT
Iba1 (Direct)
β-tubulin
GFAP
Homer1
S100B
CCP
Laminin (Direct)
SV2A
PV
SLC2A1
vGluT2
Laminin
5th staining round
DAPI
Double cortin
cFOS
COXIV
Calnexin
DDX3X
NF-H
Lamin B1 (Direct)
SST
Giantin
Necab2
Lamin B1
ABAT
Iba1 (Direct)
β-tubulin
GFAP
Homer1
S100B
CCP
Laminin (Direct)
SV2A
PV
SLC2A1
vGluT2
Laminin
SOX2 (Direct)
CALB2
MAP2
vGluT1
NeuN
Synapsin1

**Supplementary Figure 15. Cumulative visualization of the spectro-temporal unmixing result shown in Figure 6.** Scale bar = 50 μm.

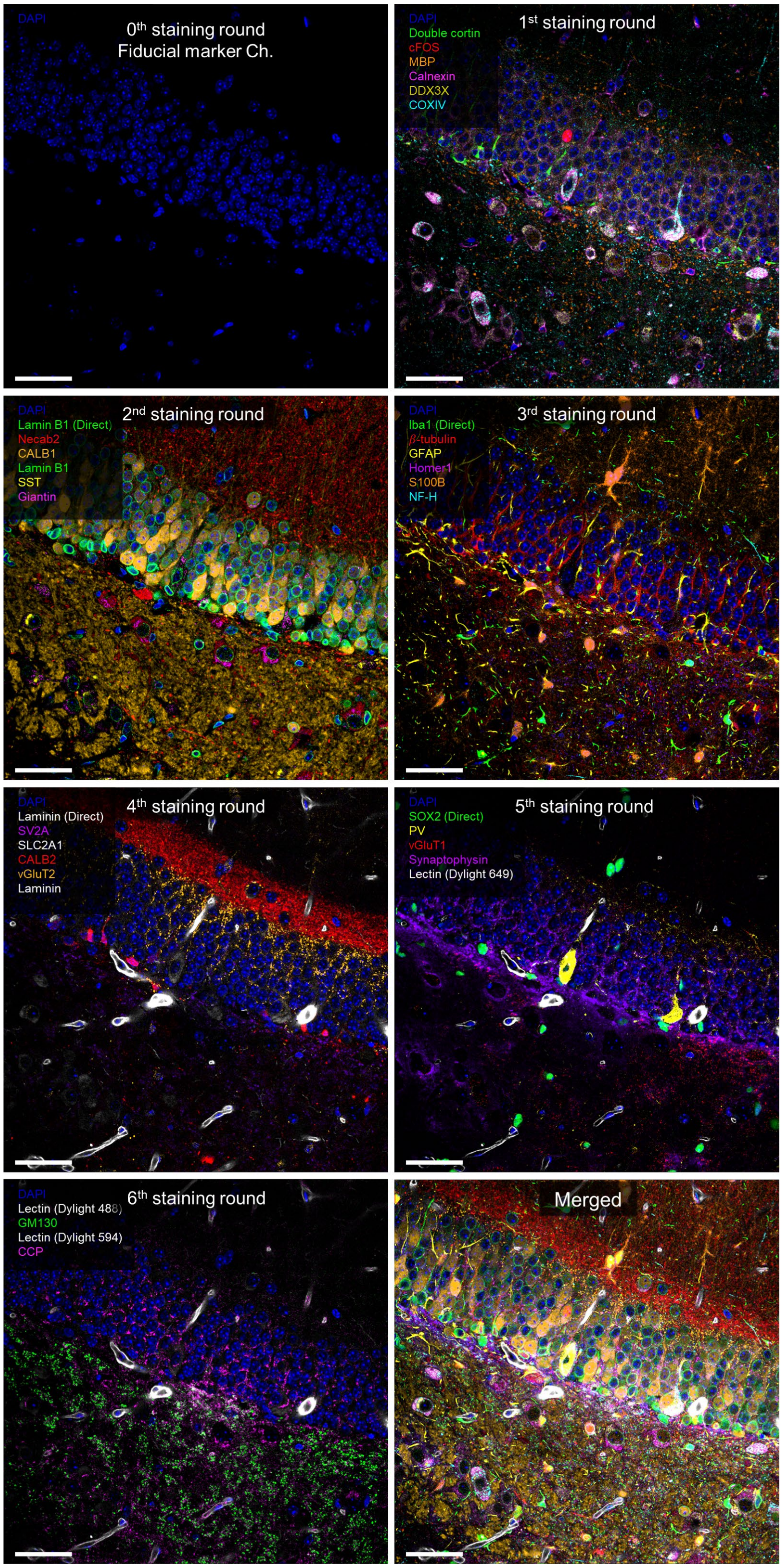


**Supplementary Figure 16. An additional demonstration of the spectro-temporal unmixing with different protein panels.** 33-plex imaging was performed in 6 rounds. Scale bar = 50 μm.

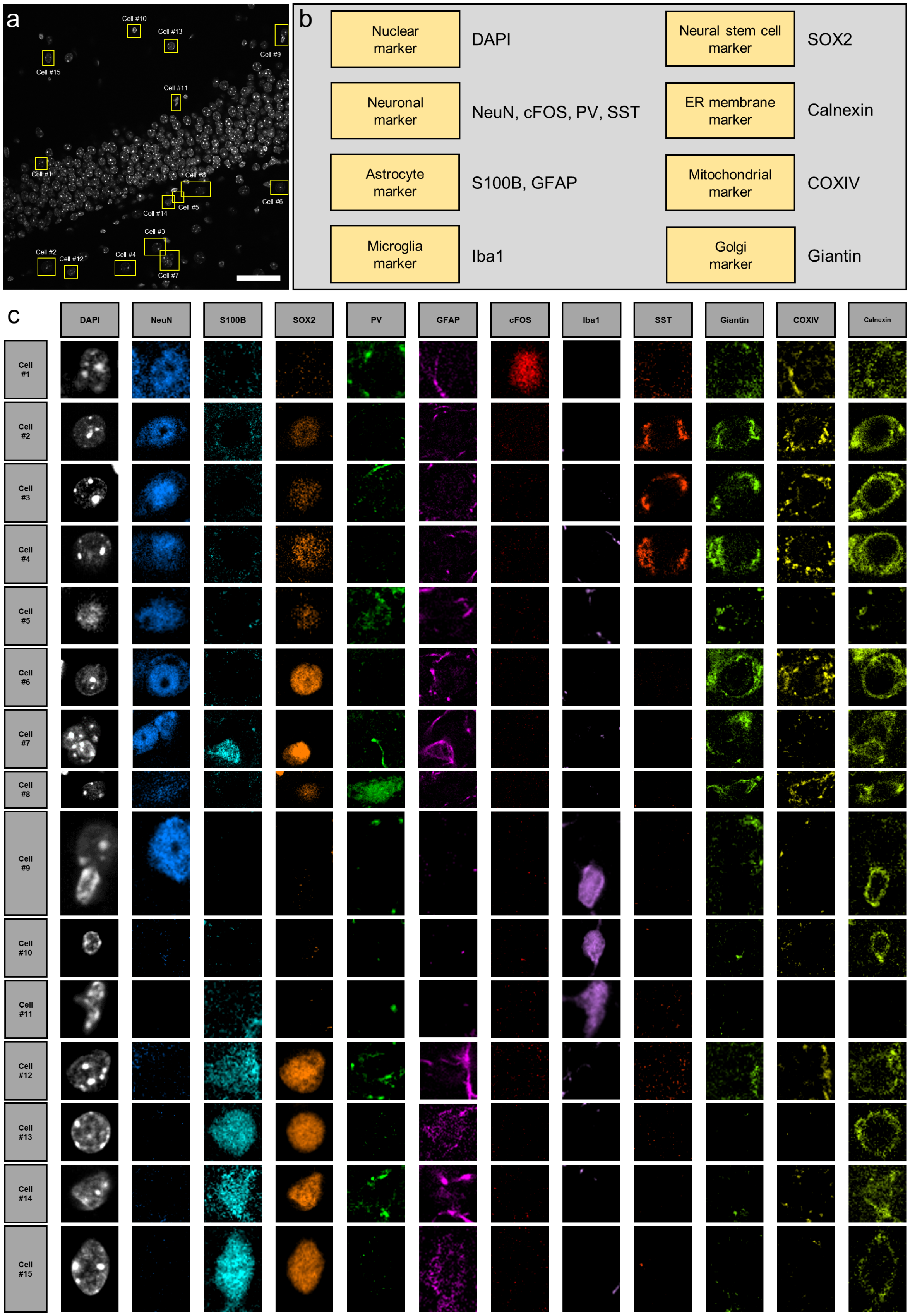
a
Cell #10
Cell #13
Cell #9
Cell #15
Cell #11
Cell #1
Cell #8
Cell #6
Cell #5
Cell #14
Cell #3
Cell #2
Cell #12
Cell #4
Cell #7
b
Nuclear marker
DAPI
Neural stem cell marker
SOX2
Neuronal marker
NeuN, cFOS, PV, SST
ER membrane marker
Calnexin
Astrocyte marker
S100B, GFAP
Mitochondrial marker
COXIV
Microglia marker
Iba1
Golgi marker
Giantin
c
DAPI
NeuN
S100B
SOX2
PV
GFAP
cFOS
Iba1
SST
Giantin
COXIV
Calnexin
Cell #1
Cell #2
Cell #3
Cell #4
Cell #5
Cell #6
Cell #7
Cell #8
Cell #9
Cell #10
Cell #11
Cell #12
Cell #13
Cell #14
Cell #15

**Supplementary Figure 17. Cells with different protein expression profiles found in Figure 6d–i.** (**a**) An image of the DAPI channel. Fifteen regions were selected, and the protein expression profiles of the cells in those regions were displayed in **c**. (**b**) List of the proteins shown in **c**. (**c**) Protein expression profiles of the cells in the boxed regions of **a**. Scale bar = 50 μm.

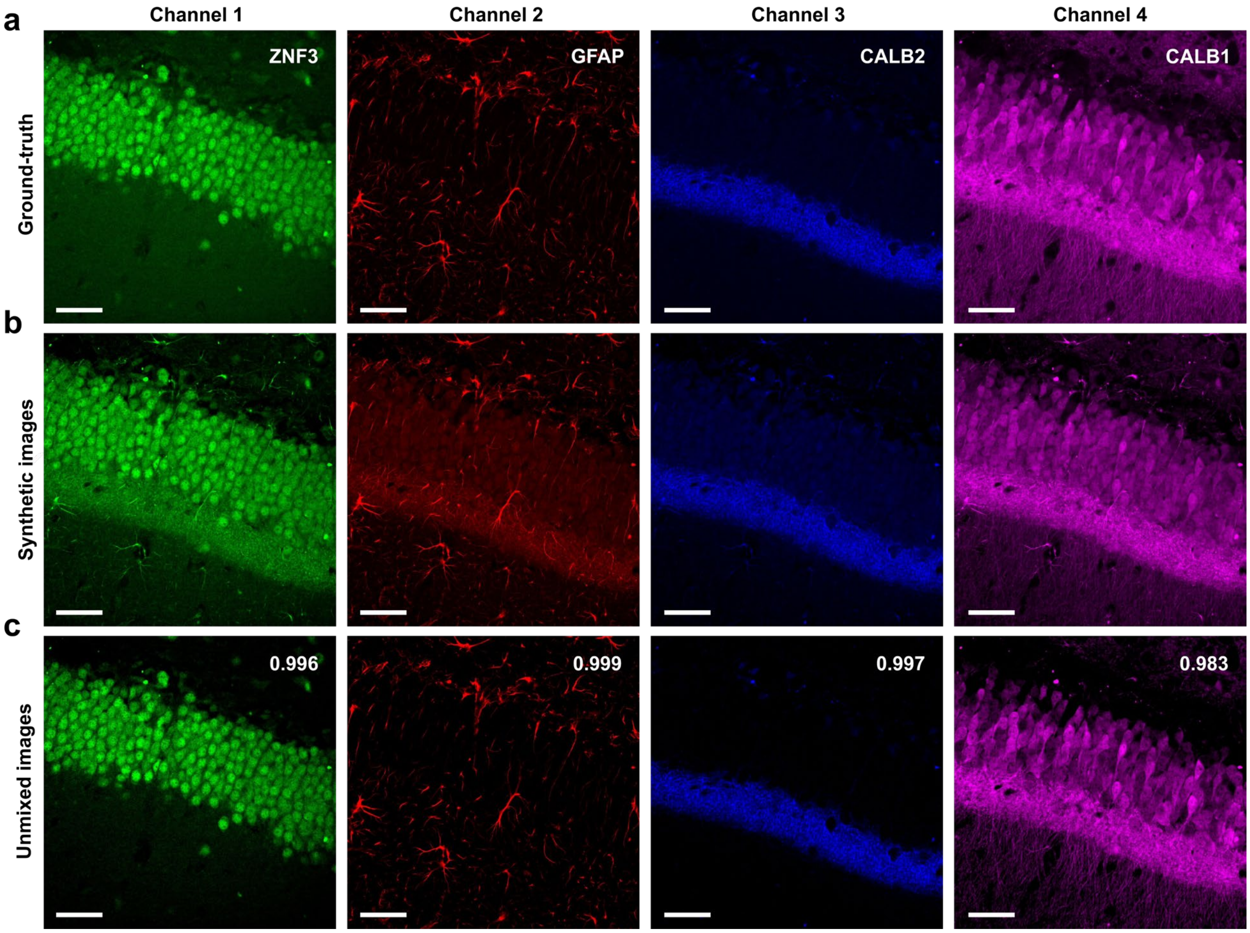


**Supplementary Figure 18. Simulation result of the four-color spectral unmixing.** (**a**) Four protein images of a mouse brain slice used for synthesizing mixed images. Green: ZNF3, red: GFAP, blue: CALB2, magenta: CALB1. (**b**) Synthetic mixed images. (**c**) Results of the unmixing of the mixed images shown in **b** via neural unmixing. The Pearson correlation coefficients between the ground truth and the unmixed images are displayed within each unmixed result. All Pearson correlation coefficient is above 0.98. Scale bar = 50 μm.

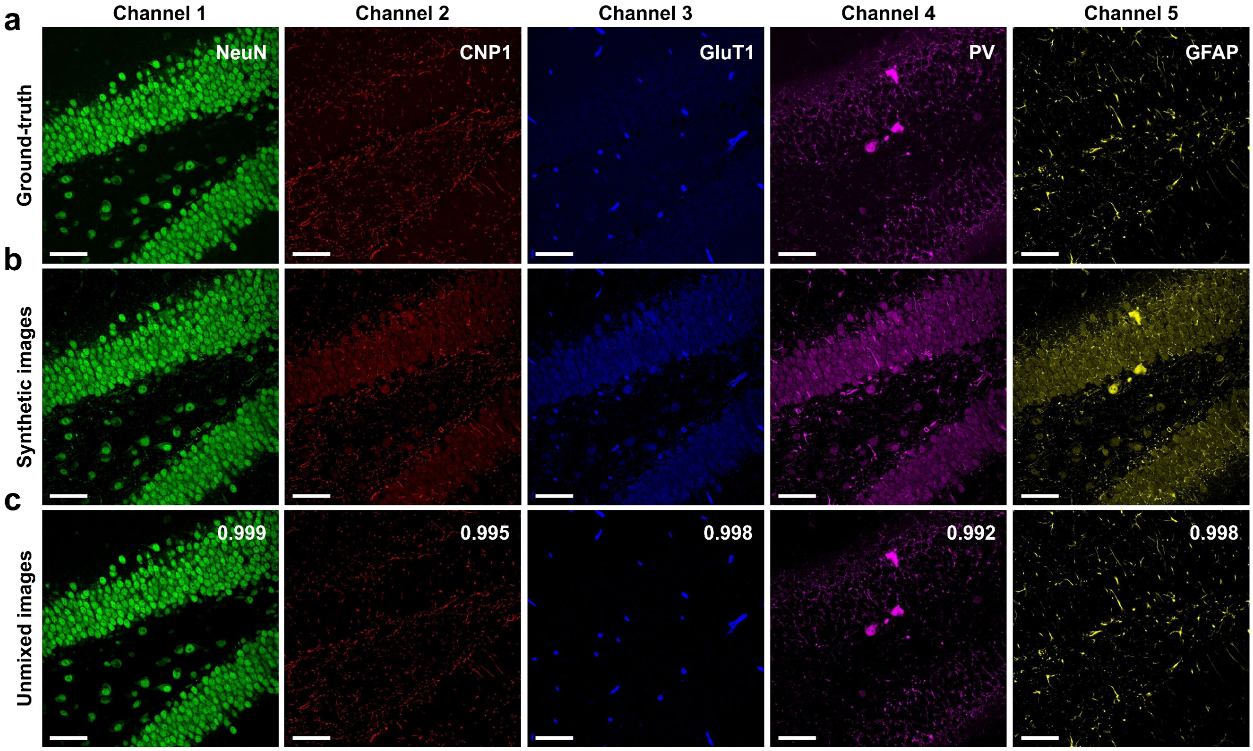


**Supplementary Figure 19. Simulation result of the five-color spectral unmixing.** (**a**) Five protein images of a mouse brain slice used for synthesizing mixed images. Green: NeuN, red: CNP1, blue: GluT1, magenta: PV, and yellow: GFAP. (**b**) Synthetic mixed images. (**c**) Results of the unmixing of the mixed images shown in **b** via neural unmixing. The Pearson correlation coefficients between the ground truth and the unmixed images are displayed within each unmixed result. All Pearson correlation coefficients are above 0.99. Scale bar = 50 μm.

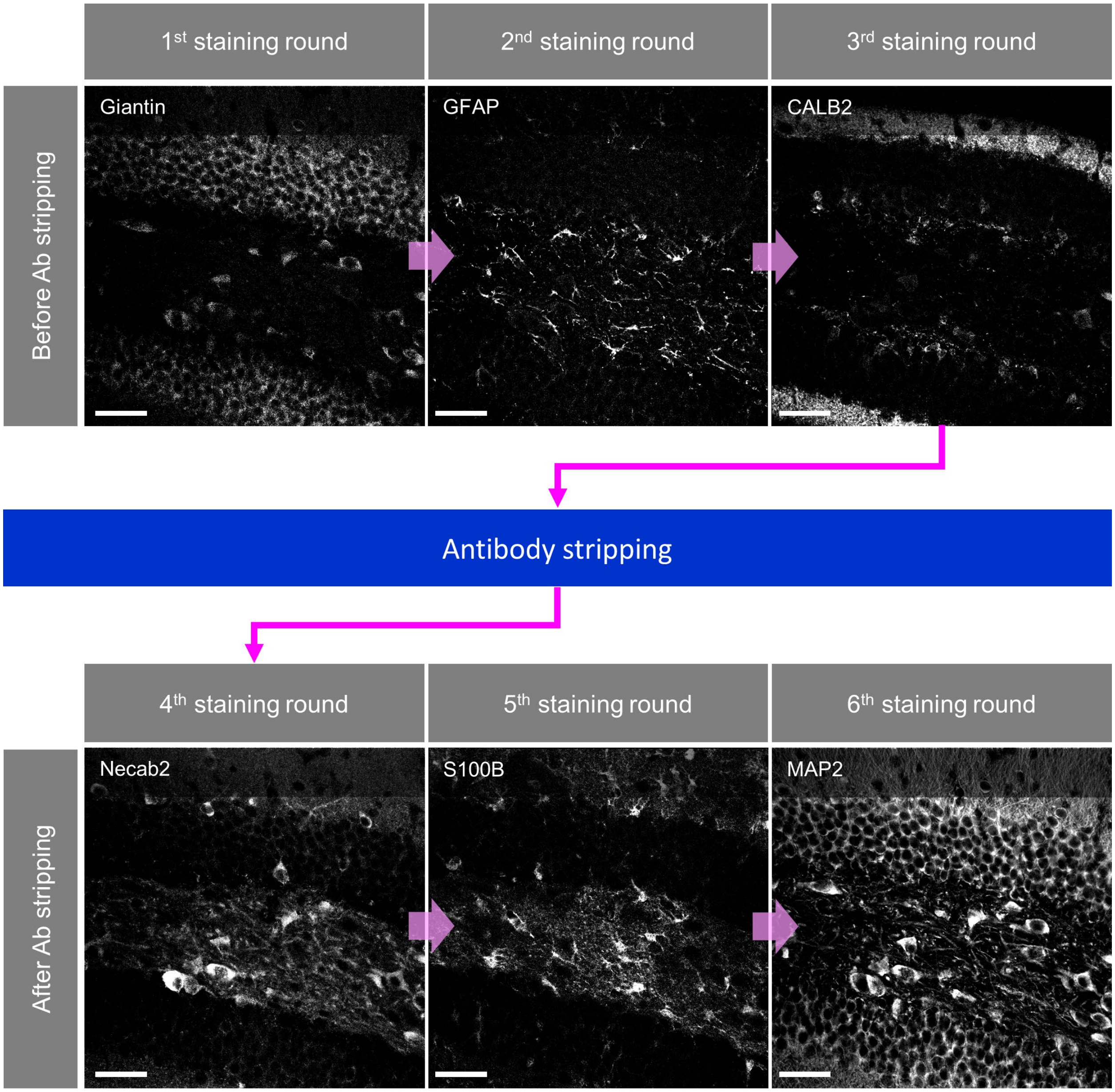


**Supplementary Figure 20. Combination of IMPASTO with an antibody stripping technique.** Three-plex imaging was performed on an FFPE mouse brain slice. Then, the antibodies were stripped by using NewBlot Nitro 5X stripping buffer. Following the antibody stripping process, another three-plex imaging was performed. Scale bar = 50 µm.

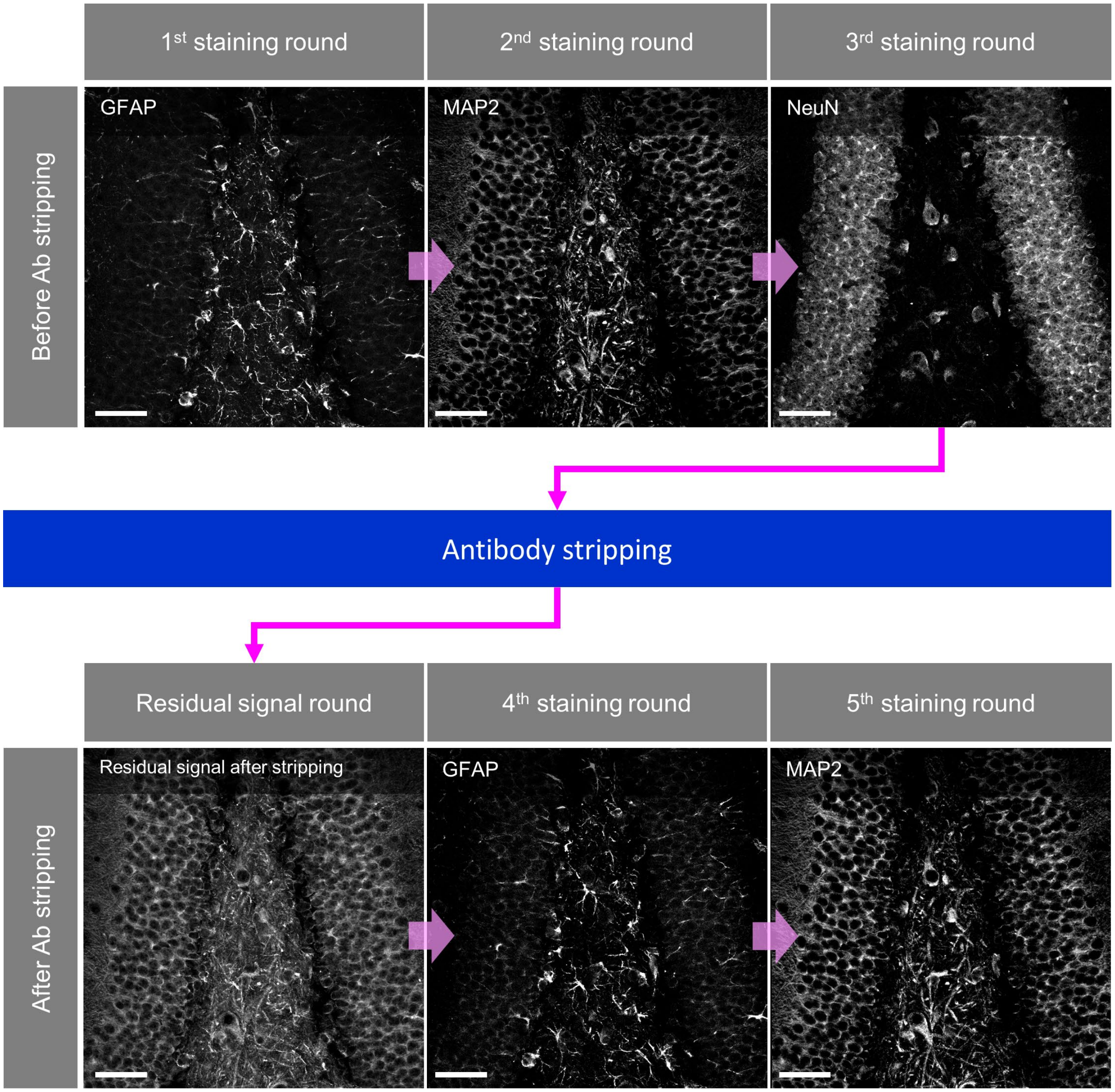


**Supplementary Figure 21. The use of IMPASTO to remove residual signals.** Three-plex imaging was performed on an FFPE mouse brain slice. Then, the antibodies were stripped by using NewBlot Nitro 5X stripping buffer. Although the antibody signals were not completely removed by the antibody stripping process, additional cyclic staining was performed. The image of the fourth protein (=GFAP) was acquired by unmixing the fourth-round image and the image of the residual signals using the neural unmixing algorithm. Scale bar = 50 μm.

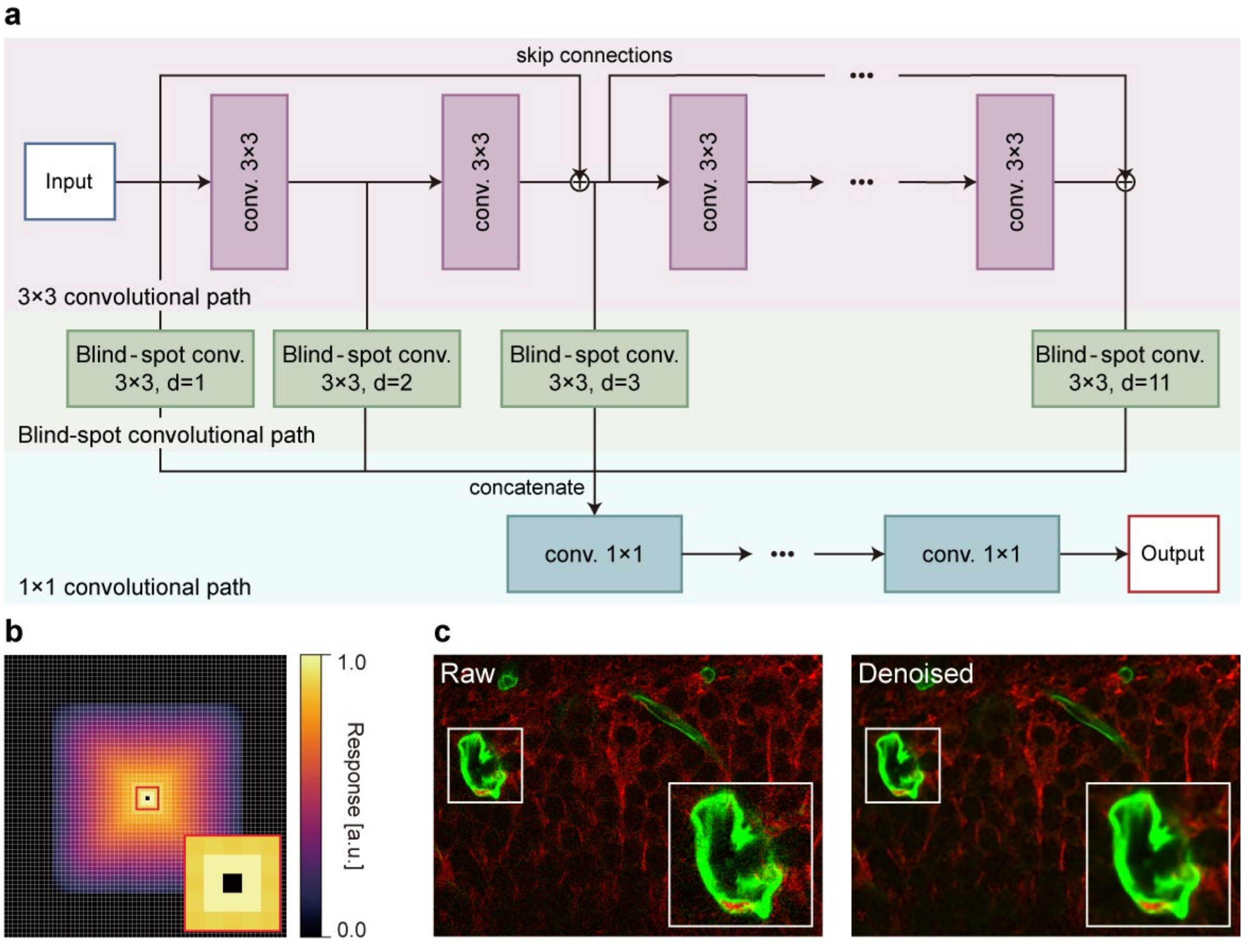


**Supplementary Figure 22. The self-supervised denoising.** (**a**) Denoising network consists of $3 \times 3$ convolutional path, blind-spot convolutional path, and $1 \times 1$ convolutional path. (**b**) The impulse response of the denoising network has a size of $43 \times 43$ pixels with a blind spot in the center. The inset in bottom right zooms in on the small red box at the center. (**c**) A noisy input image and a denoised image. The inset in bottom right zooms in on the small white box on the left.
